# Input-specific synaptic depression shapes temporal integration in mouse visual cortex

**DOI:** 10.1101/2023.01.30.526211

**Authors:** Jennifer Y. Li, Lindsey L. Glickfeld

## Abstract

Efficient sensory processing requires the nervous system to adjust to ongoing features of the environment. In primary visual cortex (V1), neuronal activity strongly depends on recent stimulus history. Existing models can explain effects of prolonged stimulus presentation, but remain insufficient for explaining effects observed after shorter durations commonly encountered under natural conditions. We investigated the mechanisms driving adaptation in response to brief (100 ms) stimuli in L2/3 V1 neurons by performing *in vivo* whole-cell recordings to measure membrane potential and synaptic inputs. We find that rapid adaptation is generated by stimulus-specific suppression of excitatory and inhibitory synaptic inputs. Targeted optogenetic experiments reveal that these synaptic effects are due to input-specific short-term depression of transmission between layers 4 and 2/3. Thus, distinct mechanisms are engaged following brief and prolonged stimulus presentation and together enable flexible control of sensory encoding across a wide range of time scales.

## Introduction

Adaptation plays a key role in dynamic regulation of sensory systems. A proposed function of sensory adaptation is to maximize stimulus information from the environment while minimizing metabolic cost of the nervous system, also known as the efficient coding hypothesis^1–3^. This optimization is particularly important in the context of naturalistic stimuli, which contain highly correlated temporal and spatial structure^4–6^. By reducing neuronal sensitivity to repeated or relatively constant stimulus features, adaptation can improve the metabolic efficiency of stimulus representation. To accomplish this, sensory systems must account for the redundancy of current stimulus features by referencing a stored memory of stimulus statistics and modulate responses accordingly.

Notably, naturalistic stimuli fluctuate over a wide range of timescales, spanning milliseconds to many minutes, and these dynamics are further enriched by self-generated movements during active sensation^7,8^. Therefore, reducing redundant encoding across timescales requires sensory systems to concurrently store stimulus statistics across a wide range of temporal contexts. Indeed, measured effects of adaptation can accrue over a variety of timescales. Responses recorded from neurons in visual, auditory, and somatosensory cortices are best predicted by sets of temporal filters that encompass multiple timescales of stimulus history^9,10^. However, whether adaptation acts to improve encoding through a singular mechanism that acts on multiple timescales, or multiple mechanisms, is still unknown.

Part of this ambiguity arises from the complexity of biological processes related to adaptation. Ion channel kinetics and short-term synaptic plasticity can often be fit with concurrent fast and slow time constants that differ by orders of magnitude^1,11–14^. However, studies that have systematically measured adaptation across multiple timescales provide strong evidence for contribution from multiple, distinct mechanisms. In the retina, fast and slow contrast adaptation modulate retinal circuitry in different ways^15^. At the level of primary visual cortex (V1), brief and prolonged presentation of the same visual stimulus produce distinct effects on neurons’ orientation tuning curves^16^. This idea extends even to human psychophysics, where duration and dynamics of adapter stimuli can determine not only the magnitude, but also specific features of perceived visual aftereffects^17,18^. Altogether, both perceptual and neural effects of adaptation are consistent with multiple mechanisms that act across different timescales.

Here, we investigated the mechanism underlying adaptation in layer 2/3 (L2/3) neurons in V1 of alert mice. L2/3 neurons in V1 undergo a profound degree of adaptation to brief stimulus presentations (0.1 s; rapid adaptation)^19,20^. Consistent with an efficient coding model, visual responses to repeated stimuli are suppressed more than responses to novel stimuli. Although adaptation could be inherited through many stages of visual processing prior to L2/3, the majority of this effect appears to originate within cortex, as neurons in both the visual thalamus (lateral geniculate nucleus; LGN) and the thalamic input layer of cortex (layer 4; L4) show very little effect of adaptation at this time scale^20,21^. Although cell-intrinsic mechanisms can explain adaptation effects with prolonged stimulus presentation^22,23^, they are insufficient for explaining rapid adaptation’s relatively brief time scale of induction as well as stimulus-selectivity. Instead, these features have largely been attributed to mechanisms involving inhibition and short-term synaptic plasticity^14,24–28^. However, the mechanisms engaged with rapid adaptation have yet to be directly tested.

Using a combination of *in vivo* and *in vitro* electrophysiological approaches, we measured the relative contribution of cell-intrinsic and synaptic mechanisms to this form of rapid adaptation. We find that adaptation with brief visual stimulus presentation does not engage significant hyperpolarization mechanisms. Instead, we find balanced a decrease in both excitatory and inhibitory synaptic inputs that can account for the decreasing in firing rate associated with rapid adaptation. Manipulations that directly activate L4, or decrease probability of release at L4 synapses, demonstrate that this site is both necessary and sufficient for rapid adaptation, and argue for a role of short-term depression at this synapse. Altogether, our results highlight a complementary role for cell-intrinsic and synaptic mechanisms in maintaining multiple time scales of sensory adaptation.

## Results

### Rapid adaptation reduces stimulus-evoked synaptic inputs

Visual responses of neurons in L2/3 of V1 are substantially reduced following even brief (0.1 s) visual stimuli^9,19,21^. This is largely a cortical phenomenon as neurons in the thalamic input layer of V1 (L4) undergo significantly less suppression than those in L2/3^20^. Thus, this rapid adaptation is likely due to a local mechanism affecting cell-intrinsic excitability of L2/3 neurons or the efficacy of their synaptic inputs. Previous work investigating cortical mechanisms of adaptation revealed that extended visual stimulus presentation (tens of seconds) evokes a cell-intrinsic hyperpolarization that accounts for decreased stimulus-evoked responses^22,23^. We first investigated whether rapid adaptation is also mediated by cell-intrinsic mechanisms by making intracellular membrane potential recordings of L2/3 V1 neurons in awake, head-fixed mice. Pairs of high-contrast, static gratings (0.1 s, 0.1 cycles per degree, 30° diameter) at the neuron’s preferred orientation were presented at a range of inter-stimulus intervals (ISIs) to measure the magnitude and time course of recovery from rapid adaptation (**Figure 1A**). Presentation of the baseline stimulus induces a decrease in firing rate (FR) in response to the test that is consistent with previous studies using calcium imaging and extracellular recordings^19,20^ (**Figure 1B**). At short ISIs, responses to the test stimulus are suppressed by nearly 40% (normalized FR [0.25 s ISI]: 0.62 ± 0.08; n = 13 cells; p < 0.001, paired t-test; **Figure 1C**) and recover with a time constant of nearly 1 second (τ = 0.82 s).

**Figure 1.**
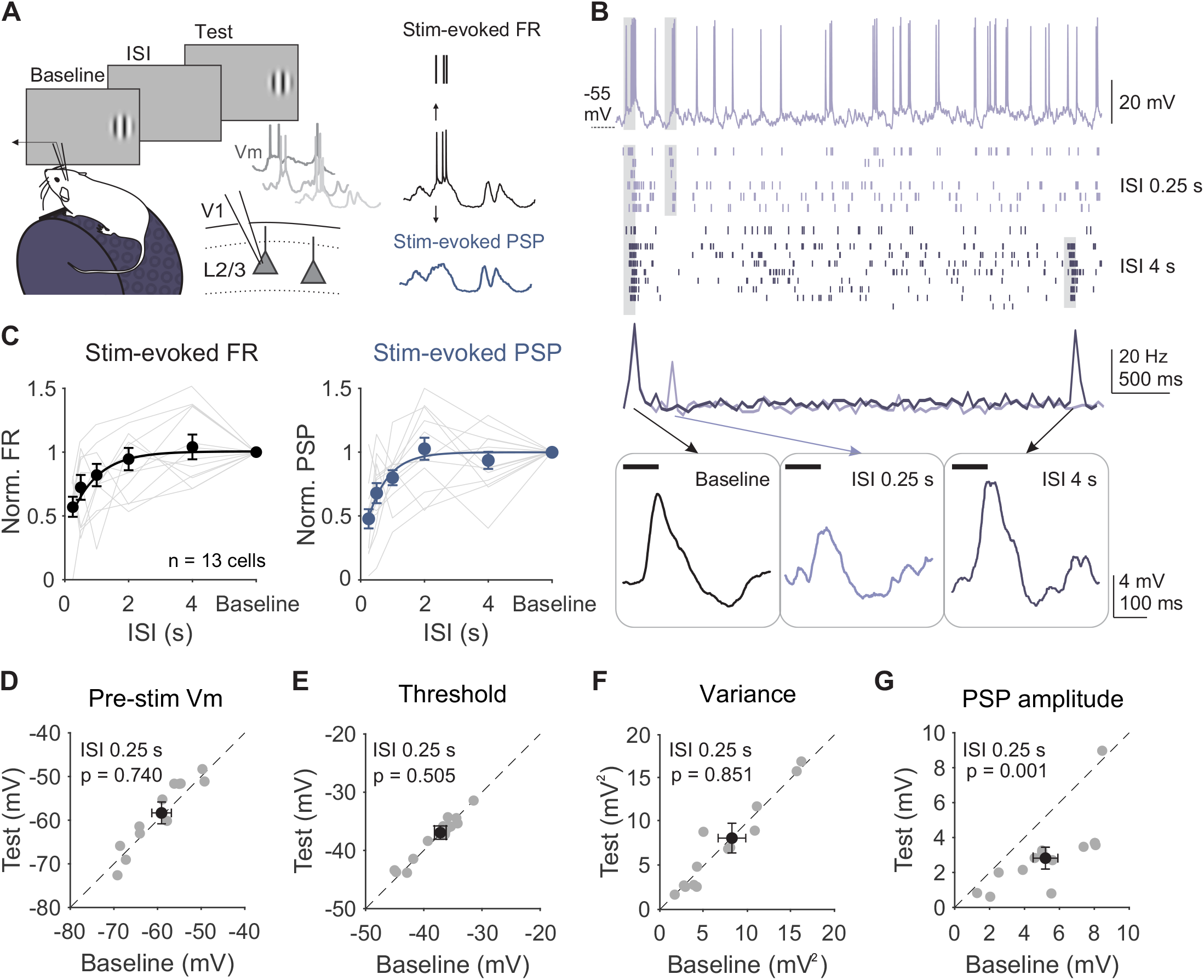
Adaptation suppresses stimulus-evoked responses in L2/3 neurons without affecting cell-intrinsic properties. **A**. Left: Recording setup and stimulus paradigm. Animals are head-fixed on a treadmill and membrane potential (Vm) of L2/3 neurons is recorded with a glass pipette. Two stimuli (baseline and test; 0.1 s) are separated by an inter-stimulus interval (ISI) varying from 0.25 to 4 s. Right: Membrane potential is separated into the stimulus-evoked firing rate (black) and stimulus-evoked post-synaptic potential (blue; PSP). **B**. Top: Membrane potential from an example cell during a 0.25 s ISI trial. Grey shading indicates stimulus presentation. Middle: Raster plot of spike output during 0.25 s (light purple) and 4 s (dark purple) ISI trials and binned peri-stimulus spike histogram (PSTH). Bottom: Subthreshold membrane potential during baseline (left) and test stimulus presentations at 0.25 s (middle) and 4 s (right) ISIs. **C**. Left: Average normalized firing rate (FR; test/baseline) as a function of ISI for individual cells (gray lines) and all cells (black circles; n = 13). Error is SEM across cells. Black line is an exponential fit (τ = 0.82 s). Right: Same as left, for average normalized PSP amplitude (τ = 0.79 s). **D**. Average membrane potential preceding baseline and test stimuli for individual cells in 0.25 s ISI trials. Black dot is mean across cells. Error bar is SEM across cells. **E-G**. Same as **D** for spike threshold (**E**), membrane variance (**F**), and PSP amplitude (**G**).

To determine whether this decrease in firing rate could be explained by a long-lasting hyperpolarization following the baseline stimulus, we compared the membrane potential preceding baseline and test stimuli. Despite the strong suppression of spike output, there is no significant hyperpolarization of membrane potential prior to the test stimulus (0.25 s ISI – baseline: -58.72 ± 2.25 mV; test: -58.25 ± 2.50 mV; p = 0.74, paired t-test; **Figure 1D**). Other properties of the recorded cell that could impact spike output are also unchanged, such as the spike threshold (0.25 s ISI – baseline: -40.07 ± 1.01 mV; test: -40.32 ± 1.15 mV; p = 0.50, paired t-test; **Figure 1E**) and membrane variance (0.25 s ISI – baseline: 7.63 ± 1.88 mV^2^; test: 7.54 ± 1.94 mV^2^; p = 0.85, paired t-test; **Figure 1F**). Additionally, most neurons have a positive correlation between the number of spikes in response to baseline and test stimuli on each trial, arguing against a cell-intrinsic fatigue effect (**Figure S1A**). Although a cell-intrinsic hyperpolarization mechanism exists in V1 L2/3 neurons, these changes in membrane potential appear only after prolonged periods of activity (**Figure S1B-D**). Instead, adaptation in response to brief stimuli greatly reduces stimulus-evoked post-synaptic potentials (PSPs), with a similar magnitude (normalized PSP [0.25 s ISI]: 0.51 ± 0.11; **Figure 1B-C, G**) and time course of recovery (τ = 0.79 s) as is seen for changes in spike output. Therefore, rapid adaptation engages a synaptic, rather than cell-intrinsic, mechanism to reduce stimulus-evoked responses to repeated stimuli.

Stimulus-evoked PSPs are generated by the sum of both excitatory and inhibitory inputs onto a post-synaptic cell. Increases in inhibition, decreases in excitation, or decreases in total conductance could all lead to reduced stimulus-evoked depolarization. To identify changes in stimulus-evoked excitation and inhibition, we made voltage clamp recordings from L2/3 neurons while presenting the same stimulus paradigm (**Figure 2A**). We recorded both excitatory currents (EPSCs) and inhibitory currents (IPSCs) from individual neurons by clamping the membrane potential near the reversal for inhibition (−70 mV) and excitation (+10 mV), respectively (n = 10 cells; **Figure 2A** and **S2A**). Consistent with our current clamp recordings, there are no changes in either the mean or the standard deviation of the holding current in the time windows preceding baseline and test stimulus onset (−70 mV: Δ current -2.33 ± 12.21 pA; p = 0.86; Δ std 0.67 ± 3.65 pA; p = 0.44; +10 mV: Δ current -2.79 ± 15.90 pA; p = 0.86, Δ std -4.14 ± 6.07 pA; p = 0.28; paired t-test for all comparisons). This argues against a role for long-lasting inhibition or changes in overall network excitability in rapid adaptation.

**Figure 2.**
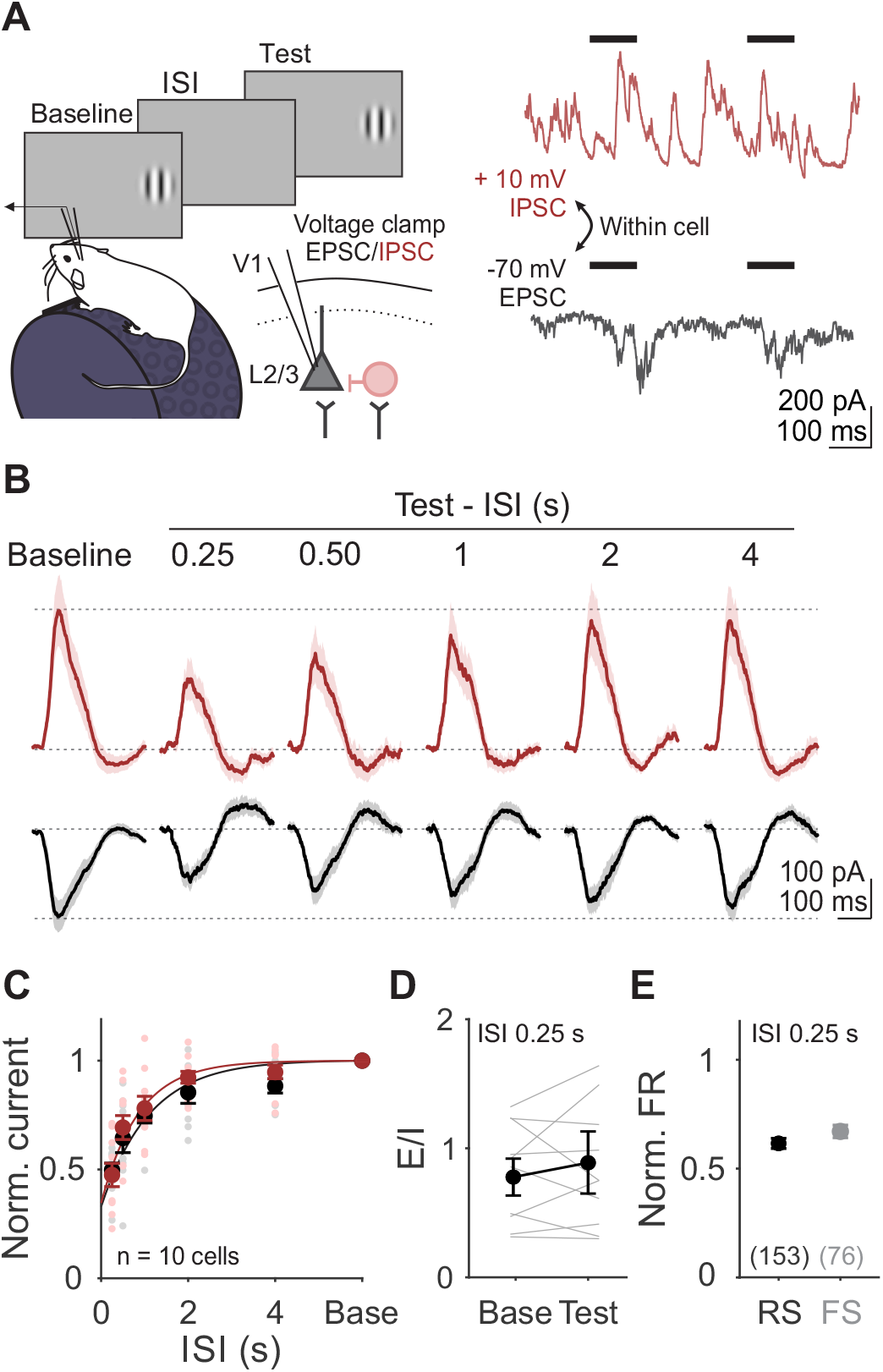
Adaptation drives a balanced reduction in stimulus-evoked excitation and inhibition. **A**. Left: Schematic of recording setup for measuring excitatory and inhibitory currents (EPSCs and IPSCs) in L2/3 neurons. Right: Single trial voltage traces from an example cell held at - 70 mV (black) and +10 mV (red), to measure EPSCs and IPSCs respectively. **B**. Grand average of stimulus-evoked EPSCs and IPSCs across all cells (n = 10) in response to baseline and test stimuli for all ISIs. Shaded error is SEM across cells. **C**. Average normalized current amplitudes (test/baseline) for EPSCs (black) and IPSCs (red) for individual cells (small dots) and across all cells (large dots). Curve is exponential fit to the average across cells for each current type. Error bar is SEM across cells. **D**. Ratio of excitation to inhibition (E/I) for the baseline and test stimulus in 0.25 s ISI trials. Grey lines are individual cells, black line is average across cells, error is SEM across cells. **E**. Comparison of visual adaptation in 0.25 s ISI trials in putative pyramidal cells (RS, black) and inhibitory interneurons (FS, gray), obtained from extracellular recordings (**Figure S3**). Error bar is SEM across units.

Instead, there is a robust decrease in the peak amplitude of stimulus-evoked excitation (normalized EPSC: 0.51 ± 0.06; p < 0.001, paired t-test; **Figure 2B-C**) and inhibition (normalized IPSC: 0.47 ± 0.09; p < 0.001, paired t-test) in response to the test stimulus relative to baseline. This reduces the overall conductance (baseline: 7.45 ± 3.11 nS; test: 3.87 ± 2.76 nS; p < 0.001, paired t-test) while preserving E/I ratio (baseline: 0.81 ± 0.16; test: 0.95 ± 0.27; p = 0.16, paired t-test; **Figure 2D**). Stimulus-evoked synaptic inputs are suppressed to a similar degree as the postsynaptic potentials (EPSC vs PSP: p = 0.98; IPSC vs PSP: p = 0.94; unpaired t-test) and firing rates (EPSC vs FR: p = 0.19; IPSC vs FR: p = 0.16; unpaired t-test) measured intracellularly, and recover at a similar time scale (τ_EPSC_ = 1.10 s; τ_IPSC_ = 0.93 s; **Figure 2C**). Thus, the decrease in synaptic drive can account for the magnitude and time course of the reduced excitability following rapid adaptation.

Notably, the magnitude and time course of changes in excitatory and inhibitory synaptic inputs are remarkably well-matched (**Figure 2C-D**). This suggests that the two may be yoked by a shared mechanism, such as a decrease in the excitation onto both excitatory and inhibitory cells. Indeed, we find a comparable decrease in the firing rate of both putative excitatory (regular-spiking [RS], n = 135 units) and inhibitory (fast-spiking [FS], n = 67 units) neurons in L2/3 (normalized FR [0.25 s ISI]: RS = 0.65 ± 0.02, FS = 0.71 ± 0.04, p = 0.08, unpaired t-test; **Figures 2E and S3A-B**). Altogether, these observations are consistent with short-term depression of excitatory synapses onto both excitatory and inhibitory neurons.

### Adaptation acts at specific excitatory synapses

If reduction in excitation and inhibition in L2/3 neurons *in vivo* is generated by short-term synaptic depression of intracortical synapses, changes in synaptic inputs should reflect the features of this type of plasticity. First, we expect that repeated visual stimulus presentations will drive increasing depression of visual responses and eventually saturate at a level determined by the balance between time constants of vesicle depletion and replenishment^11,12,14^. Second, these effects should be restricted to the specific subset of synapses activated by features of the baseline stimulus (**Figure 3A**). To test these predictions, we measured EPSCs and IPSCs in response to static gratings of matched and orthogonal orientations (0.1 s duration each). We presented five stimuli of the same orientation (baseline and test 1-4) to measure accumulation/saturation, followed by an orthogonal grating to measure specificity (test 5).

**Figure 3.**
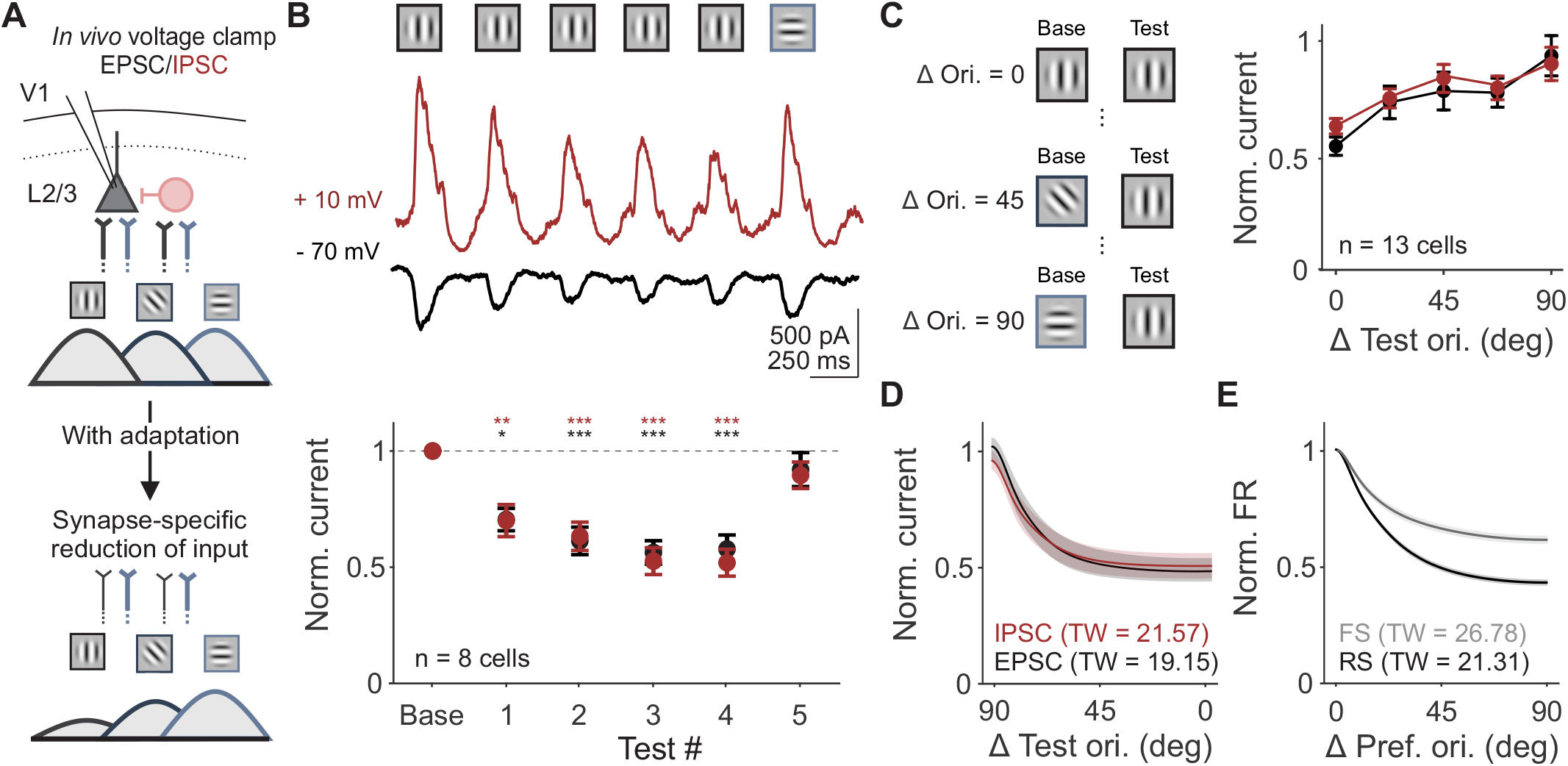
Changes in synaptic input are selective to previously activated synapses. **A**. Schematic of proposed model of synapse-specific effect of adaptation on excitatory inputs from L4 to L2/3. This generates orientation-selective decrease of synaptic inputs to both excitatory and inhibitory L2/3 neurons. Color of axons correspond to L4 inputs to L2/3 synapses tuned to vertical (black) versus horizontal (blue) orientations. Line thickness represents strength of inputs. **B**. Top: Visual stimulus paradigm with repeated presentation of the same stimulus orientation (baseline and test 1-4) followed by an orthogonal orientation (test 5). Middle: Average stimulus evoked EPSCs (black) and IPSCs (red) for an example cell. Bottom: Average normalized current (test/baseline) for all cells (n = 8). Response to the orthogonal orientation is normalized to its own baseline. Error bar is SEM across cells. **C**. Left: Schematic of stimuli presented to measure the tuning width of adaptation. Test orientation was kept constant while the baseline orientation varied. Right: Average normalized current (test/baseline, where the baseline is the same orientation as the test) as a function of similarity between baseline and test stimuli for EPSCs and IPSCs for all cells (n = 13). **D**. Average adaptation tuning curve fits from data in **C**. Shaded error is SEM across cells. Tuning width (TW) is half-width at half-max. **E**. Average orientation tuning curve fits from extracellular recording of V1 RS (black) and FS (gray) units.

Our results confirm both predictions. First, we find that suppression of both EPSCs and IPSCs accumulate and saturate over the five repeated stimuli (n = 8 cells; test 1 vs baseline 1, p = 0.04 for EPSCs and p = 0.009 for IPSCs; test 2-4 vs baseline, p < 0.001 for EPSCs and IPSCs; all comparisons within test 2-4, p > 0.05 for EPSCS and IPSCS; one-way ANOVA with post hoc Tukey test; **Figure 3B**). Second, excitation and inhibition evoked by the fifth, orthogonal test stimulus are not significantly different from the baseline response at that orientation (test 5 vs baseline: EPSCs p = 0.89; IPSCs p = 0.98; paired t-test), consistent with a synapse-specific mechanism.

Across all stimuli presented, excitation and inhibition remain balanced relative to baseline levels (p = 0.81; effect of current type, two-way ANOVA), which we attribute to a parallel decrease in excitatory drive to pyramidal cells and interneurons. We further tested this by probing the orientation selectivity of adaptation of excitation and inhibition. If decreases in EPSCs and IPSCs are the result of short-term depression at excitatory synapses, the orientation selectivity of adaptation of excitation and inhibition should be matched. Additionally, this selectivity should reflect the tuning of spike output in pyramidal neurons, which are generally more narrowly tuned than interneurons^29–31^. To measure the tuning width of adaptation, we measured excitation and inhibition in response to pairs of stimuli with orientation differences between 0 and 90 degrees, sampled in 22.5 degree increments (0.25 s ISI only; **Figure 3C**). We find that the degree of adaptation depends on orientation difference (n = 13 cells; two-way ANOVA: main effect of orientation, p = 0.009) but not current type. EPSCs and IPSCs undergo a similar degree of suppression across all orientation differences (main effect of current type: p = 0.32).

To determine whether the orientation selectivity of this suppression matches the orientation tuning of spike output in V1 neurons, we fit individual neurons’ normalized EPSCs and IPSCs in response to the test stimulus with a von Mises function (**Figure 3D**). We then compared these intracellular adaptation tuning curves to the orientation tuning curves of either RS or FS units obtained in extracellular recordings (**Figure 3E** and **S3C**). We find that the bandwidth of adaptation observed in EPSCs and IPSCs more closely matches the bandwidth of orientation tuning of RS units than FS units (tuning width (TW): RS = 21.31 ± 1.25, FS = 26.78 ± 1.91; EPSC = 19.15 ± 3.36, IPSC = 21.57 ± 3.44). The match between the orientation-selectivity of adaptation of IPSCs and RS tuning further supports the idea that changes in excitation and inhibition are yoked by a shared short-term depression mechanism that reduces excitation onto both classes of L2/3 neurons.

### Activation of L4 depresses excitatory inputs in L2/3

Our data suggest that rapid adaptation is due to short-term depression of excitatory synapses in L2/3. If so, direct activation of synaptic inputs onto L2/3 neurons should induce short-term depression and be sufficient to mimic the effects of visual adaptation. To test this prediction, we optogenetically activated inputs to L2/3 *in vitro* in slices from mice expressing Channelrhodopsin-2 (ChR2) selectively in L4 pyramidal neurons (Scnn1a-Tg3-Cre x Ai32 mice) by targeting the blue excitation light to L4 below the recorded cell (**Figure 4A**; **STAR Methods**). Optogenetic activation of cells in L4 for 0.1 s activates monosynaptic and polysynaptic excitatory inputs to L2/3 neurons (**Figure 4B**). Repeated stimulation of L4 (baseline and test), reveals a history-dependent reduction of these optogenetically-evoked EPSCs. As with the *in vivo* recordings, at short ISIs responses to the test stimulus are suppressed by nearly 40% (normalized EPSC amplitude [0.25 s ISI]: 0.63 ± 0.01; n = 11 cells; p <0.001, paired t-test; **Figure 4C**) and responses recover with a time constant of nearly 1 second (τ = 1.03 s). Therefore, the long-lasting suppression of excitatory input to L2/3 neurons observed with rapid adaptation *in vivo* can be reproduced by engaging a local, activity-dependent mechanism in V1.

**Figure 4.**
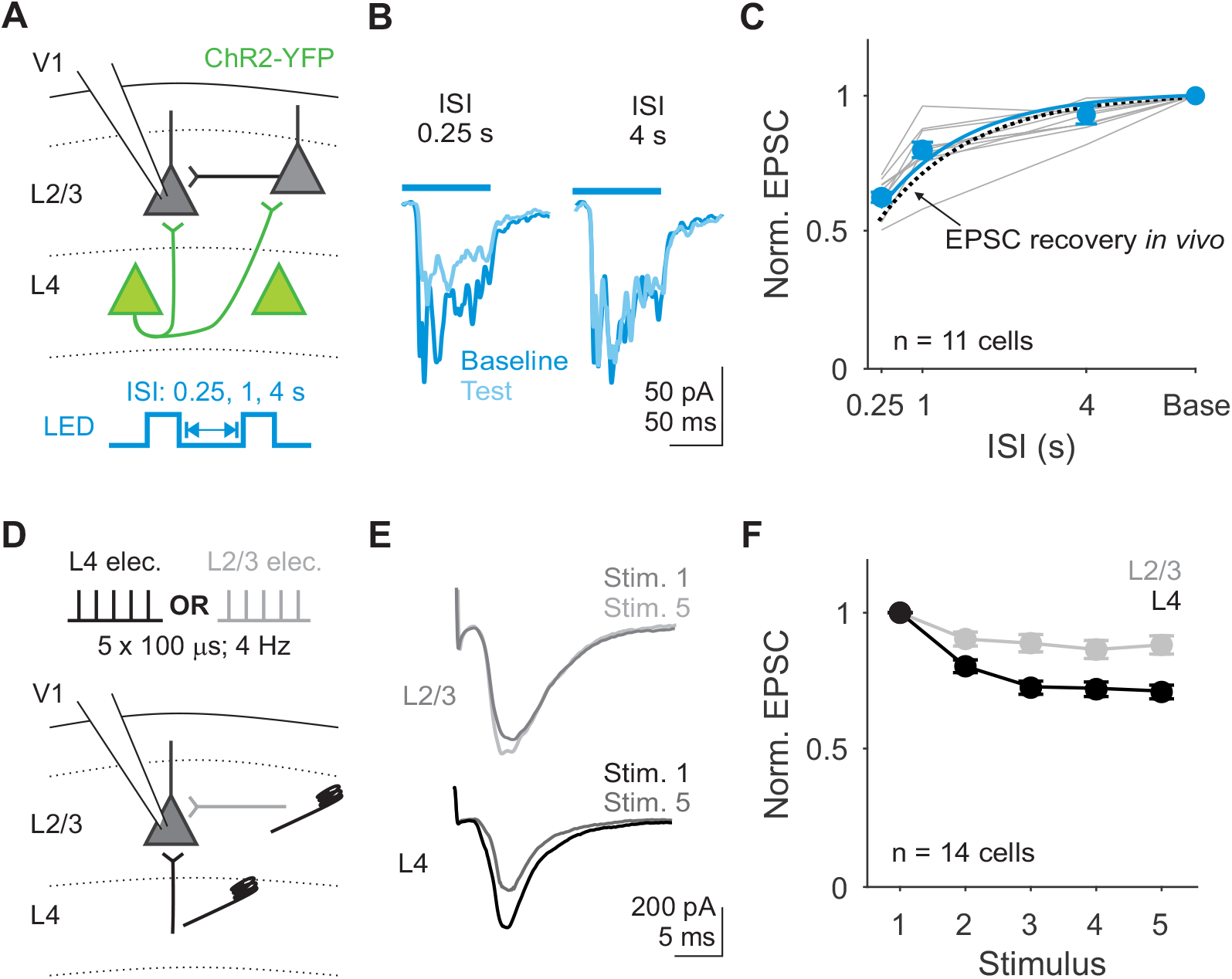
Excitatory inputs to L2/3 neurons decrease with repeated stimulation *in vitro*. **A**.Schematic of setup for recording EPSCs in L2/3 neurons during optogenetic stimulation of L4. Two 0.1 s square pulses of blue light (baseline and test) were used to activate L4 neurons. **B**. Average traces during baseline (dark blue) and test (light blue) stimuli from an example cell during 0.25 s (left) versus 4 s (right) ISI trials. **C**. Average normalized EPSC amplitudes (test/baseline) as a function of ISI for each cell (gray) and the across all cells (blue). Blue line is exponential fit to the average across cells. Dashed line is exponential fit from EPSCs recorded *in vivo* in **Figure 2**. Error bar is SEM across cells (n = 11). **D**. Schematic for recording EPSCs from a L2/3 pyramidal cell while electrically stimulating L4 or L2/3 inputs on alternating trials. **E**. Average EPSCs from an example cell in response to stimulation of L2/3 (top; gray) or L4 (bottom; black). **F**. Average EPSC amplitudes normalized to the first stimulus in response to L2/3 (gray) and L4 (black) stimulation. Error bar is SEM across cells.

In the context of the local V1 circuit, L4 stimulation *in vitro* could drive short-term depression at L4 to L2/3 synapses or at L2/3 to L2/3 synapses. To determine whether these synapses depress equally or in an input-specific manner^32,33^, we used electrical stimulation to selectively drive monosynaptic inputs from L4 or L2/3 onto L2/3 neurons (**Figure 4D**; **STAR Methods**). Repeated electrical stimulation of L4 *in vitro* is sufficient to depress EPSCs recorded in L2/3 (L4: P2/P1 = 0.82 ± 0.02, P5/P1 = 0.75 ± 0.03, n = 14 cells; **Figure 4E**). Although L4 electrical stimulation could also activate non-L4 axons passing through L4, direct optogenetic activation of L4 neurons at the same frequency depresses EPSCs to a similar extent (**Figure S4**). In contrast, L2/3 excitatory inputs onto the same cells depress significantly less (P2/P1 = 0.91 ± 0.02, P5/P1 = 0.88 ± 0.04; two-way ANOVA: main effect of input layer, p < 0.001; **Figure 4F**). Thus, adaptation in V1 L2/3 neurons likely arises from short-term depression at specific excitatory synapses originating from L4 neurons.

### Activation of L4, but not L2/3, is sufficient to drive adaptation in vivo

To test whether activation of L4 is sufficient to drive adaptation *in vivo*, we made extracellular recordings from transgenic mice expressing ChR2 in L4 (**Figure 5A**). Units were identified as L4 or L2/3 neurons based on waveform position relative to layer boundaries determined by the visually-evoked current source density (**Figure S5**). In agreement with the *in vitro* results, 0.1 s of repeated optogenetic activation of L4 neurons significantly decreases responses in L2/3 neurons, but not L4 neurons (**Figure S6A-C**). In order to investigate the interaction between optogenetic activation and subsequent visually driven responses, we used a 0.5 s sinusoidal light stimulus (**Figure S6D-F**). Trials were randomly interleaved to present visual stimulation alone or visual stimulation preceded by optogenetic stimulation of L4 (**Figure 5B**). We then compared adaptation induced by visual stimuli under control conditions (Test_control_/Baseline_control_: “Visual adapt”; **Figure 5C**) and adaptation induced by optogenetic stimulation (Baseline_opto_/Baseline_control_: “Opto. adapt” **Figure 5C**).

**Figure 5.**
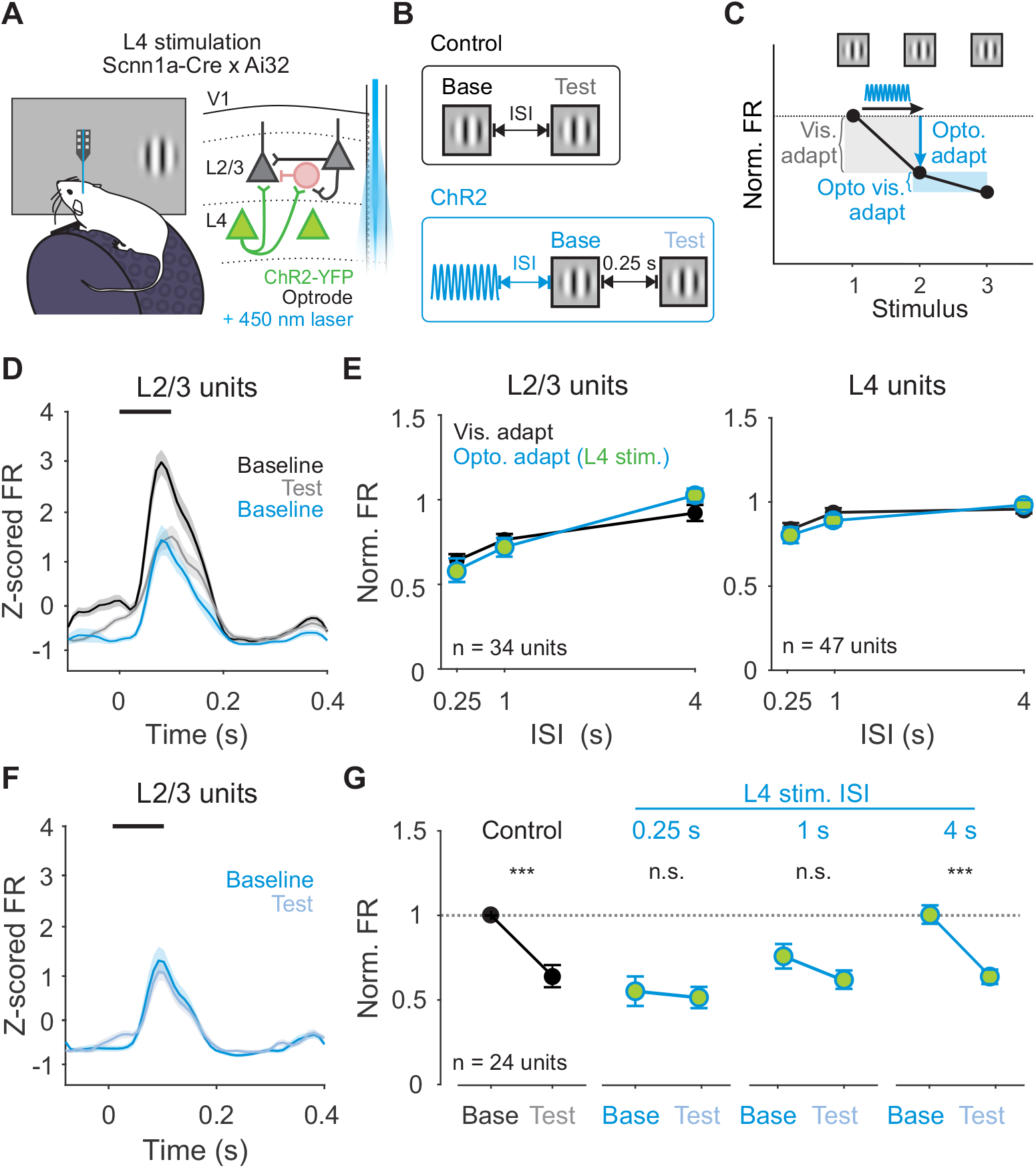
Activation of L4 neurons is sufficient to recapitulate the effects of visual adaptation. **A**. Schematic of in vivo extracellular recording setup with optrode coupled to a 450 nm laser. **B**. Structure of control trials (black) and ChR2 activation trials (blue). On control trials, baseline and test stimuli are presented with varying ISI. On ChR2 activation trials, 0.5 s of sinusoidal blue light is used to activate L4 neurons optogenetically at varying intervals prior to baseline visual stimulus presentation. **C**. Visual adaptation is quantified as the response to the test divided by the response to the baseline stimulus (gray shaded box). Optogenetic adaptation is quantified as the response to the baseline stimulus in ChR2 activation trials divided by the response to the baseline stimulus in control trials (blue arrow). Optogenetic visual adaptation is quantified as the response to the test stimulus divided by the baseline stimulus following on ChR2 activation trials (blue shaded box). **D**. Average z-scored PSTH for L2/3 units during baseline (black) and test (gray) stimuli in control trials and baseline stimulus in ChR2 activation trials (blue; n = 34 units). Black line indicates stimulus presentation. Shaded error is SEM across unit. **E**. Comparison of visual adaptation (black) and optogenetic adaptation (blue) in L2/3 (left) and L4 (right) units. Green fill indicates optogenetic stimulation of L4. **F**. Average z-scored PSTH for L2/3 units during baseline (blue) and test (light blue) stimuli in L4 ChR2 activation trials. **G**. Visual adaptation (black) and Optogenetic visual adaptation (blue) with 0.25 s ISI at increasing intervals after L4 stimulation (0.25 s, 1 s, 4 s). Normalized firing rate is calculated relative to baseline visual response in control trials (horizontal dashed line). Error bar is SEM across units.

Consistent with previous work, on control trials V1 neurons in L2/3 are suppressed by visual adaptation at short ISIs (L2/3 Visual adapt [0.25 s ISI]: 0.64 ± 0.04; n = 34 cells; **Figure 5D-E**) while neurons in L4 undergo significantly less adaptation^20^ (L4 Visual adapt [0.25 s ISI]: 0.82 ± 0.04; n = 47 cells; p < 0.001, unpaired t-test). Optogenetic activation of L4 neurons generates effects similar to visual adaptation in both L2/3 and L4: baseline visual responses are more strongly reduced in L2/3 (L2/3 Opto. adapt [0.25 s ISI]: 0.60 ± 0.07; n = 34 units; **Figure 5D-E**) than in L4 (L4 Opto. adapt [0.25 s ISI]: 0.81 ± 0.05; n = 47 units; p < 0.001, unpaired t-test). The time scale of recovery from optogenetic adaptation is also similar to recovery from visual adaptation. Across all ISIs, optogenetic adaptation is indistinguishable from visually evoked adaptation (two-way ANOVA, effect of stimulation type: L2/3 p = 0.93, L4 p = 0.47). Notably, in a subset of L2/3 neurons that are not activated by L4 optogenetic stimulation, visual responses are unaffected even shortly after ChR2 activation (L2/3 Opto. adapt [0.25 s ISI]: laser active neurons [n = 26 units] vs not laser active neurons [n = 8 units], p < 0.001; **Figure S6G**). L4 neurons showed a similar, but not significant trend (L4 Opto. adapt [0.25 s ISI]: laser active neurons [n = 39 units] vs not laser active neurons [n = 8 units], p = 0.11; un-paired t-test; **Figure S6H**). Thus, activation of L4 is sufficient to reproduce the magnitude, recovery, and layer-specific effects of visual adaptation.

Although optogenetic stimulation of L4 is sufficient to drive adaptation, it is possible that similar effects are produced through a non-overlapping, parallel mechanism to visual adaption. Because the effects of visual adaptation saturate quickly with additional stimulus presentations (**Figure 3B, 4F** and **5C**), we reasoned that if optogenetic stimulation and visual adaptation act through the same mechanism, stimulation of L4 should also reduce subsequent visual adaptation^20^. Conversely, persistence of strong visual adaptation would indicate engagement of distinct mechanisms. To test this, we compared the magnitude of visual adaptation at short (0.25 s) ISIs in control trials versus after optogenetic stimulation (Test_opto_/Baseline_opto_: “Opto. visual adapt”; (**Figure 5C**). We find that following optogenetic stimulation of L4, responses to the test stimulus show little effect of visual adaptation (L2/3 Test_opto_ vs Baseline_opto_ [0.25 s ISI]: n = 24 units; p = 0.49). Consequently, visual adaptation in L2/3 is significantly reduced following optogenetic adaptation (L2/3 Visual adapt vs Opto. visual adapt [0.25 s ISI] p < 0.001, paired t-test); **Figure 5F-G**). The occlusion of adaptation in L2/3 by stimulation of L4 indicates that adaptation evoked by visual and optogenetic stimulation likely act through the same mechanism.

While optogenetic stimulation of L4 is sufficient to induce visual adaptation in L2/3, this stimulation also activates recurrent and feedback inputs within L2/3, which could generate the effects we observe. To test whether activation of L4 is necessary for driving visual adaptation, we used *in utero* electroporation to selectively express ChR2 in L2/3 pyramidal cells and made extracellular recordings under the same experimental conditions (**Figure 6A-B**). Firing rates in L2/3 are not reduced following L2/3 stimulation (Opto. adapt [0.25 s ISI]: 1.05 ± 0.06; n = 27 units; p = 0.94, paired t-test; **Figure 6C-D**), and the magnitude of this effect is significantly smaller than occurs in response to both visually evoked adaptation (p < 0.001; paired t-test) and L4 stimulation (p = 0.007; unpaired t-test). In addition, unlike L4, L2/3 stimulation does not occlude visual adaptation (Opto. visual adapt [0.25 s ISI]: 0.68 ± 0.08; n = 27 units; Test_opto_ vs Baseline_opto_ [0.25 s ISI]: p = 0.002, paired t-test; **Figure 6E-F**). Overall, our results are consistent with the preferential short-term depression at L4 inputs to L2/3 observed *in vitro*. We find that activation of L4, but not L2/3, can recapitulate the effects of visual adaptation. Thus, activation of the L4 to L2/3 synapse is both necessary and sufficient for visual adaptation in L2/3.

**Figure 6.**
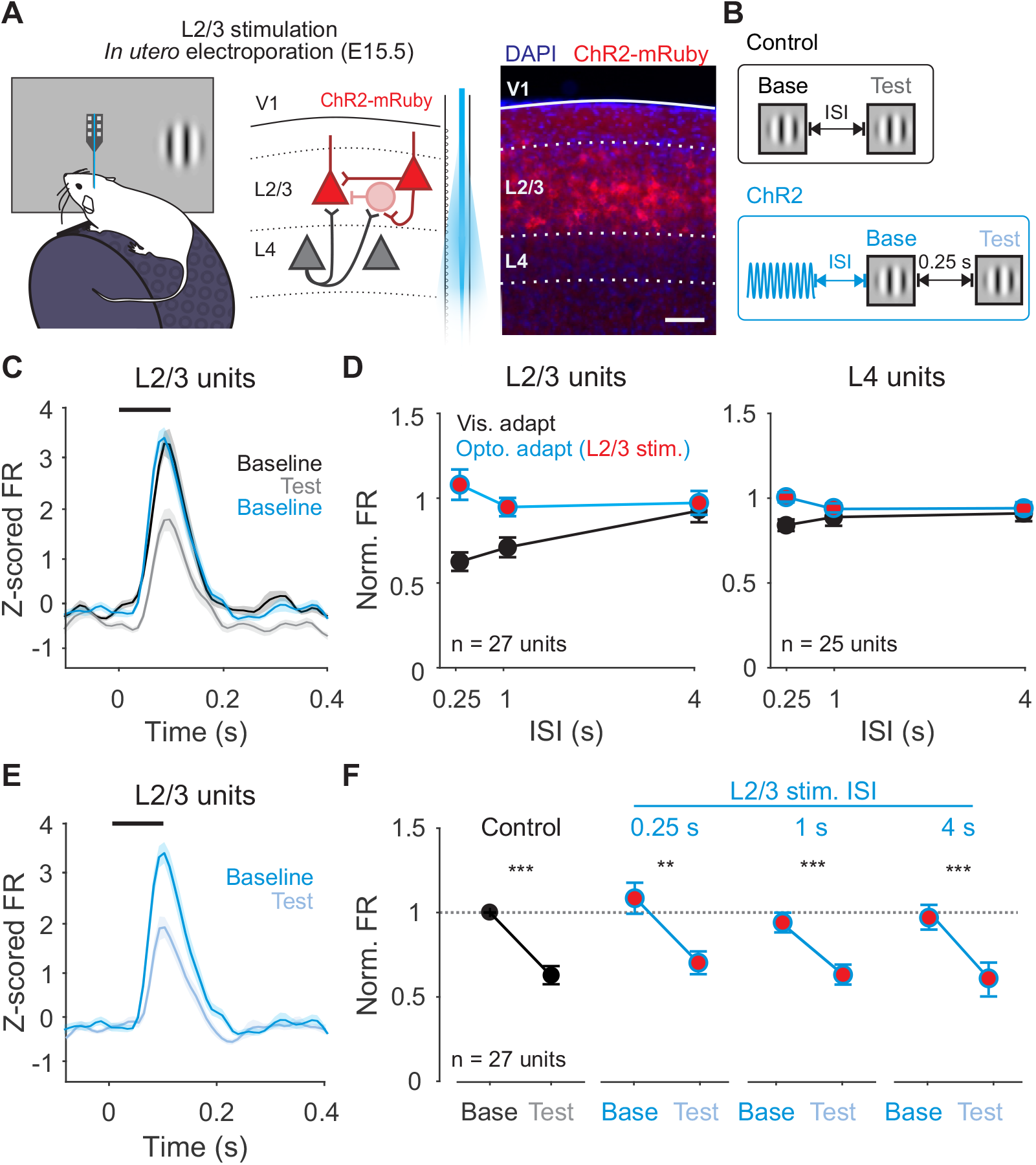
Activation of L2/3 neurons does not recapitulate the effects of visual adaptation. **A**. Left: Schematic of in vivo extracellular recording setup in mice expressing ChR2 in L2/3 neurons. Right: expression of ChR2-mRuby in L2/3 neurons following in utero electroporation. Scale bar is 100 μm. **B**. Structure of control trials (black) and ChR2 activation trials (blue). **C**. Average z-scored PSTH for L2/3 units during baseline (black) and test (gray) stimuli in control trials and baseline stimulus in ChR2 activation trials (blue; n = 27 units). Black line indicates stimulus presentation. Shaded error is SEM across units. **D**. Comparison of visual adaptation (black) and optogenetic adaptation (blue) in L2/3 (left) and L4 (right) units. Red fill indicates optogenetic stimulation of L2/3. **E**. Average z-scored PSTH for L2/3 units during baseline (blue) and test (light blue) stimuli in L2/3 ChR2 activation trials. **F**. Visual adaptation (black) and Optogenetic visual adaptation (blue) with 0.25 s ISI at increasing intervals after L2/3 stimulation (0.25 s, 1 s, 4 s). Normalized firing rate is calculated relative to baseline visual response in control trials (horizontal dashed line). Error bar is SEM across units.

### Rapid adaptation results from short-term depression at L4 to L2/3 synapses

Short-term depression is associated with activity-dependent depletion of readily releasable vesicles at high release probability (Pr) synapses^11^. To test whether short-term depression at L4 synapses is necessary for rapid adaptation, we optogenetically manipulated Pr using the modified mosquito opsin, eOPN3, which enables reversible inhibition of vesicle release^34^. With green light exposure, eOPN3 activates a G_i/o_ pathway to inhibit calcium channels and SNARE complex formation, reducing vesicle release and decreasing depletion^11,34^. Thus, we can use eOPN3 to decrease Pr selectively at L4 synapses and test whether this also decreases short-term depression and rapid adaptation.

We expressed eOPN3 in L4 neurons by injecting a Cre-dependent viral construct in Scnn1a-Tg3-Cre mice and confirmed its effects using *in vitro* whole-cell recordings of EPSCs in L2/3 neurons (**Figure 7A**). We used a small spot of green light positioned over the recorded cell to activate eOPN3 expressed at L4 axon terminals and measured EPSCs evoked with a 4 Hz train of electrical stimulation in L4 (**Figure 7B**). To ensure the effects were specific to eOPN3 activation in L4 axons, on alternating trials we recorded EPSCs evoked by placing a second stimulation electrode in L2/3, ∼100 μm from the recorded cell to avoid the ascending L4 axons. Due to the relatively slow off-kinetics of eOPN3, we performed these experiments in a block-wise structure (**Figure 7A**). After a block of control trials, the eOPN3 block was initiated with 10 s of green light exposure with an additional 0.5 s of green light exposure preceding each trial in the block. We then returned to control conditions to measure the time course of recovery.

**Figure 7.**
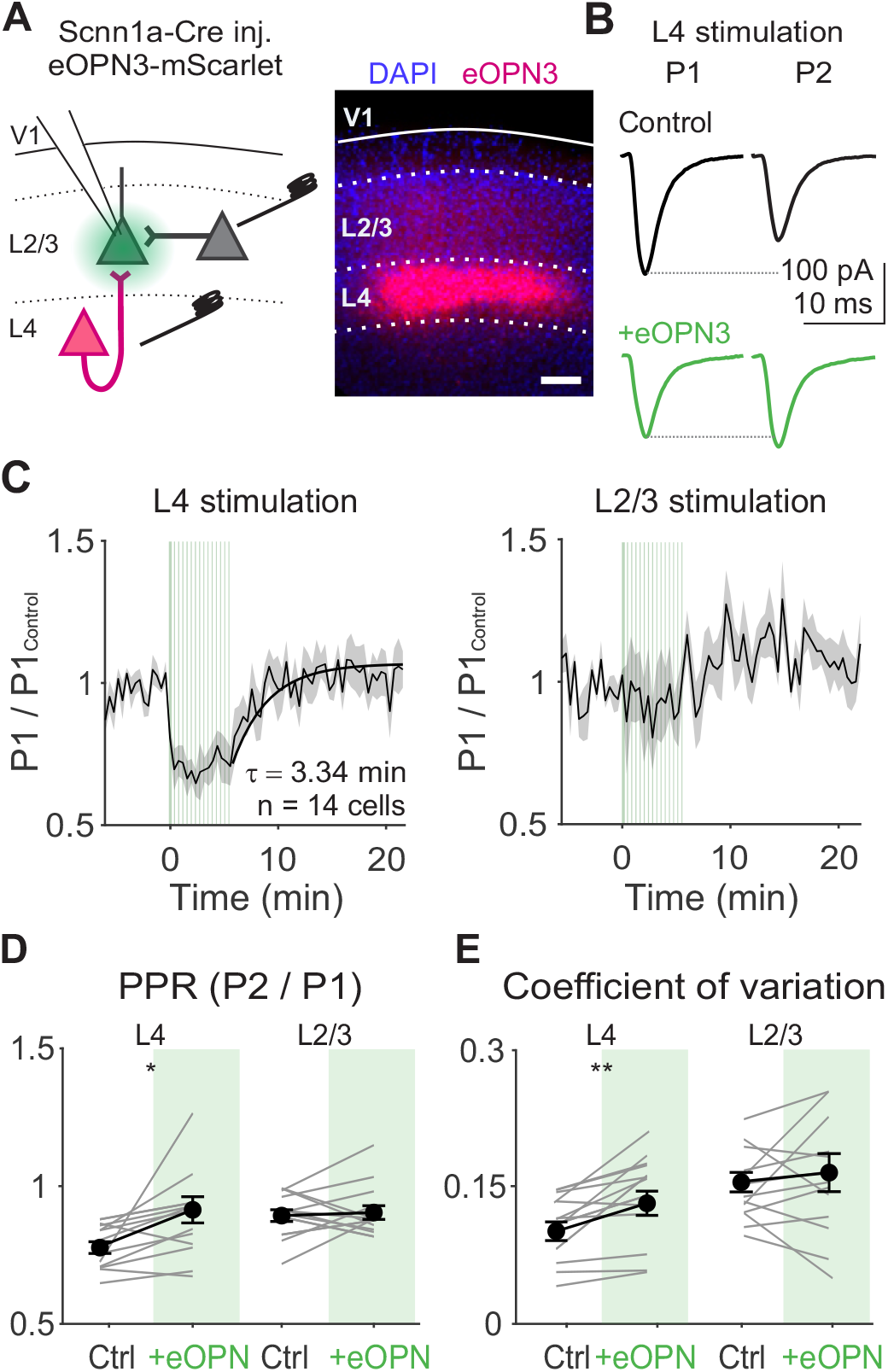
Activation of eOPN3 in L4 terminals reduces probability of release at inputs onto L2/3 neurons. **A**. Left: Schematic of *in vitro* recording setup for recording EPSCs in L2/3 neurons while electrically stimulating L4 or L2/3. eOPN3 expressed in L4 neurons is activated with green light over L4 terminals in L2/3. Right: Example image of viral expression pattern. Scale bar is 100 μm. **B**. EPSCs from an example cell in response to L4 stimulation during first (P1) and second (P2) stimuli in a train (4 Hz), either before (black) or after eOPN3 activation (green). **C**. Average time course of normalized P1 EPSC amplitudes following L4 (left) or L2/3 (right) stimulation aligned to the time of eOPN3 activation (n = 14 cells). Vertical green lines indicate eOPN3 activation trials: induction of 10 s of pulsed green light prior to visual stimulus presentation, followed by a top-up of 0.5 s of pulsed green light prior. Black curve is exponential fit to recovery. Shaded error is SEM across cells. **D**. Paired pulse ratio (PPR) during L4 or L2/3 stimulation for individual cells (gray lines) and the average of all cells (black) in control (white) and after eOPN activation (green). Error bar is SEM across cells. **E**. Same as **D**, for coefficient of variation.

Consistent with a reduction in Pr, activation of eOPN3 significantly reduces the amplitude of EPSCs elicited by L4 electrical stimulation (**Figure 7C;** P1_eOPN3_/P1_Baseline_: 0.63 ± 0.03, p < 0.001, paired t-test), increases the paired-pulse ratio (p = 0.02; **Figure 7D**), and increases the coefficient of variation (p = 0.02; **Figure 7E**). In contrast, EPSCs evoked by L2/3 electrical stimulation are significantly less suppressed than L4 stimulation (P1_eOPN3_/P1_Baseline_: 0.89 ± 0.18, L4 vs L2/3: p = 0.003, paired-t-test; **Figure 7C**) and have no significant change in paired-pulse ratio (p = 0.69, paired t-test; **Figure 7D**) or coefficient of variation (p = 0.32; **Figure 7E**). Following the eOPN3 activation block, the amplitude of evoked L4 EPSCs recover over a few minutes (τ = 3.34 min). The reversible and selective nature of the suppression suggests an effect on vesicle release, rather than unrelated instabilities during recording. Thus, optogenetic inhibition of L4 terminals can reduce short-term depression and vesicle depletion in a pathway-specific manner.

We next determined whether decreasing Pr and short-term depression at L4 synapses prevents rapid adaptation of visual responses *in vivo*. To test this, we recorded V1 neurons extracellularly while presenting pairs of static gratings (0.25 s ISI), and activated eOPN3 using the same block-wise paradigm as we validated *in vitro*, illuminating L4 axons in V1 with green light via an optic fiber outside the brain (**Figure 8A-B**). To quantify the effect of this manipulation on visual responses, we compared responses to the baseline stimulus on control and eOPN3 activation trials (**Figure 8C** and **S7A**). This manipulation produces a range of effects on stimulus-evoked firing rates: neurons with less than a 20% change in firing rate were categorized as stable, while neurons that decreased or increased by more than 20% as inhibited or facilitated, respectively (**Figure 8D**). Consistent with our manipulation largely targeting L4 to L2/3 synapses, most neurons in L2/3 are inhibited following eOPN3 activation (inhibited: 67/105; stable: 28/105; p < 0.001, Chi-squared test; **Figure 8E**), whereas most neurons in L4 are stable (inhibited: 18/61; stable: 34/61; p = 0.003, Chi-squared test;).

**Figure 8.**
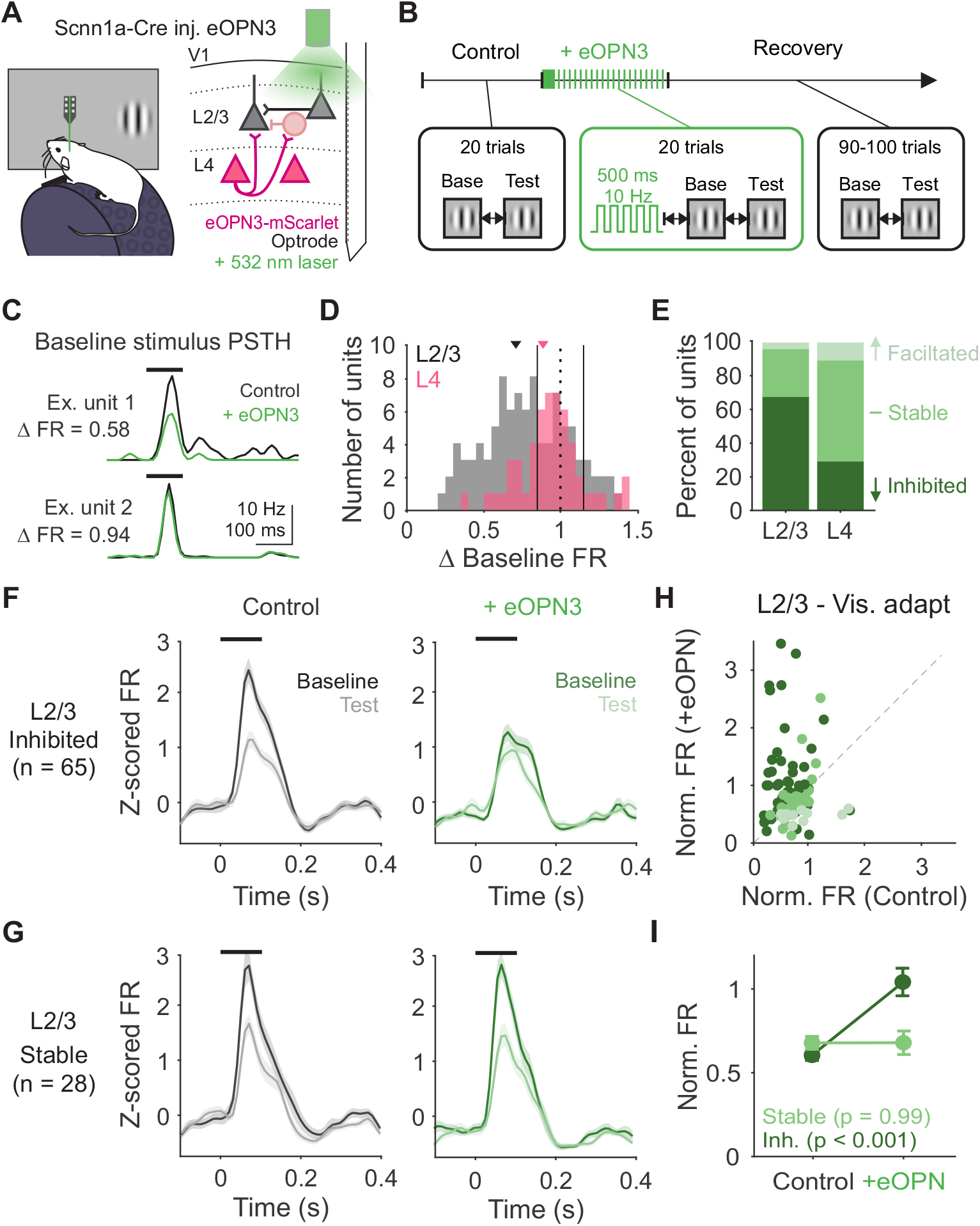
Decreasing probability of release at L4 terminals decreases visual adaptation *in vivo*. **A**. Schematic of recording setup and eOPN3 expression with green light illumination outside of the brain. **B**. Block-wise trial structure for measuring effects of eOPN3 activation on visual adaptation. Visual stimuli are always presented with 0.25 s ISI. eOPN3 activation block consist of an induction of 10 s of pulsed green light prior at the start of the block, followed by a top-up of 0.5 s of pulsed green light prior to visual stimulus presentation on each trial. **C**. PSTHs for two example units in control (black) and eOPN3 activation (green) trials. Δ FR is calculated as the change in peak stimulus-evoked response. **D**. Distribution of change in visually-evoked responses to the baseline stimulus in L4 (pink; n = 61) and L2/3 (gray; n = 105) units. Vertical solid lines indicate thresholds for categorization as inhibited (< 0.8), stable (> 0.8 and < 1.2), or facilitated (> 1.2). **E**. Percent of units categorized as inhibited, stable, or facilitated in L2/3 and L4. **F**. Average z-scored PSTH of inhibited L2/3 units (n = 65) in response to baseline (dark) and test (light) stimuli during control trials (left, black) and during eOPN3 activation trials (right, green). Black line indicates stimulus presentation. Shaded error is SEM across units. **G**. Same as **F**, for stable L2/3 units (n = 28). **H**. Comparison of normalized response (test/baseline) in control and eOPN3 activation trials, for all L2/3 units colored by categorization in **E. I**. Average normalized response for inhibited (dark green) and stable (light green) units in L2/3. Error bar is SEM across units.

If suppression of neurons in L2/3 is indicative of decreased Pr at L4 inputs to those neurons, visual adaptation should be most affected in neurons L2/3 neurons inhibited by eOPN3 activation. Indeed, inhibited neurons in L2/3 undergo significantly less visual adaptation after eOPN3 activation (p < 0.001, paired t-test; **Figure 8F, H-I**). In comparison, there is no change in the adaptation of stable neurons in L2/3 (p = 0.99; **Figure 8G-I**), inhibited neurons in L4 (p = 0.21; **Figure S7B-C**), or stable neurons in L4 (p = 0.45). These effects cannot be explained by non-specific effects of the laser, as green light activation of L4 neurons expressing only a fluorophore has no significant effect on visually-evoked firing rates of L2/3 neurons (n = 35 cells; response to baseline stimulus-p = 0.75; paired t-test; **Figure S7D-E**) or the degree of adaptation (p = 0.34). Nor could these effects be explained solely by reduced visual responses in L2/3 induced by eOPN3, as L2/3 neurons exhibit a comparable reduction of visually-evoked firing with a decrease in stimulus contrast (40% vs 80% contrast: normalized FR-0.68 ± 0.06; n = 29 cells; p < 0.001; paired t-test; **Figure S8**) with no significant effect on adaptation (p = 0.45; two-way ANOVA, effect of baseline contrast). Together, our results indicate that short-term depression at high Pr L4 to L2/3 synapses in V1 is necessary for the effects of visual adaptation.

## Discussion

We have shown that synaptic depression at feedforward synapses within primary visual cortex can explain stimulus-specific adaptation of visually-evoked responses. Our results demonstrate that features of rapidly changing visual stimuli are stored at the level of synapses through activity-dependent modulation of synaptic efficacy. Moreover, the effects of this modulation reduce sensitivity to repeated stimulus features, potentially serving to improve efficiency of stimulus encoding.

### Direct evidence for a synaptic depression mechanism in adaptation

Short-term synaptic plasticity is a fundamental feature of the nervous system that can transform physically static synapses into dynamic filters of presynaptic activity^11,35,36^. Previous *in vitro* results from electrical stimulation^12^, two-photon optogenetic input mapping^32^, and paired recordings^33,37^ have found that short-term depression is the dominant form of plasticity at L4 to L2/3 synapses in V1. In contrast to cell-intrinsic mechanisms involved at long timescales of continuous visual experience, the effects of synaptic depression can be engaged with brief, transient stimulation. Using whole-cell recordings of L2/3 neurons *in vitro* and *in vivo* we demonstrate that changes in synaptic inputs from L4 can explain the long-lasting and stimulus-specific nature of rapid adaptation of spike output.

Many studies have hinted that synaptic depression plays a role in generating adaptation to repeated stimulus presentations *in vivo*^10,24,25,27,38–43^. In cat visual cortex, repeated electrical stimulation of LGN neurons produces suppression of excitation and inhibition in cortical neurons^27^. Similarly, balanced reduction of excitation and inhibition has been observed during adaptation to whisker stimulation in barrel cortex and clicks in auditory cortex^24,38^. Although these findings are consistent with synaptic depression, these experiments did not directly test the role of short-term plasticity. In this study, we leverage cell-type specific *in vitro* and *in vivo* optogenetic manipulations to directly manipulate synaptic transmission at L4 neurons and found that activation of L4 inputs to L2/3 is both necessary and sufficient for producing the effects of visual adaptation. Notably, while we find that the majority of adaptation can be accounted for by depression at this cortical synapse, synaptic depression has been reported at both retinogeniculate and thalamocortical synapses^42,44–47^. These discrepancies could originate from differences in spontaneous activity that depend on state (awake vs anesthetized) and preparation (*in vivo* vs *in vitro*). Spontaneous thalamic activity depends on the type and depth of anesthesia, and will therefore modulate the degree of depression at thalamocortical synapses. Similarly, higher overall levels of spontaneous activity *in vivo* shift these synapses closer to saturated levels of depression at rest compared to *in vitro*^44,48,49^. Thus, the degree of adaptation along the visual hierarchy is not a fixed property of these synapses, but instead strongly depends on brain state. Our results therefore provide insight to relevant mechanisms that govern visual processing in the alert animal.

### Rapid adaptation is not associated with increased inhibition

Another mechanism that has been proposed to mediate stimulus-specific adaptation is increased inhibition. One model for increased inhibition proposes that it arises via differential synaptic plasticity at excitatory synapses from pyramidal cells to inhibitory interneurons, or from inhibitory synapses from interneurons to pyramidal cells^50–54^. In particular, facilitation of excitatory inputs onto somatostatin-expressing (SOM) interneurons is thought to sensitize them to repeated or prolonged stimulus presentations^55–57^. Indeed, manipulation of SOM interneurons selectively affects responses to frequent, but not rare stimuli in visual and auditory cortex^26,58,59^.

Contrary to this model, our recordings indicate that the adapter stimulus does not generate long-lasting inhibition in L2/3 neurons, nor does inhibition increase in response to the test stimulus. Instead, the magnitude of excitation and inhibition are tightly linked across stimulus conditions and undergo similar degrees of adaptation. The most straightforward explanation for this balanced decrease of excitation and inhibition is through a single effect of short-term depression of excitatory L4 to L2/3 synapses onto both cell types. This model is further supported by the orientation specificity of adaptation of excitation and inhibition which more closely matches the tuning of excitatory than of inhibitory neurons. Moreover, we find that FS interneurons undergo a similar degree of adaptation as neighboring RS cells. Thus, adaptation of inhibition is likely driven by short-term depression of the excitatory inputs onto L2/3 interneurons rather than short-term dynamics of their output inhibitory synapses. Notably, *in vitro* recordings reveal a strong degree of short-term depression at these inhibitory synapses^50,60–62^. Thus, it is surprising that there is no clear contribution of short-term plasticity at this synapse to driving additional adaptation of inhibition. We propose that the high firing rates of interneurons *in vivo* may put their synapses in a tonically depressed state, rendering them stable across a range of stimulus intervals^44,63^.

It is likely that our whole-cell recordings *in vivo* have limited space clamp and therefore may be underestimating the contribution of dendritic inhibition. However, we saw no dependence of the degree of adaptation of either excitatory or inhibitory currents on series resistance (**Figure S2**), arguing against a role for facilitating dendritic inhibition. Instead, the observed decrease in total synaptic input is sufficient to explain the changes observed in spike output, rendering increased inhibition unlikely to explain adaptation at this time scale of induction and recovery.

### Distinct time scales and perceptual effects of adaptation

A short-term depression mechanism predicts a distinct set of computational capacities compared to cell-intrinsic fatigue. At any moment, a single neuron’s response is determined by the sum of thousands of synaptic inputs, meaning that independent gain changes at each of these inputs can greatly increase possible modifications of activity with adaptation^64–66^. Modeling studies predict that short-term depression normalizes the strength of individual inputs to each afferent’s mean firing level to maintain postsynaptic sensitivity to changes in presynaptic firing^36,66^. Our results indicate that adaptation selectively regulates L4 to L2/3 inputs, a key cortical, feedforward synapse in visual processing. Input-specific depression at L4 but not L2/3 inputs to L2/3 neurons could shift the relative balance of information flow from feedforward to recurrent connections. Further, the cortical site of adaptation (as opposed to at the thalamocortical synapse) allows for adaptation to be orientation-specific. The stimulus specificity of short-term depression can also be extended to other forms of cortically-computed stimulus selectivity (e.g. phase or spatial frequency) to reduce redundant encoding across multiple features^67,68^. Synaptic depression has also long been proposed to act as a low-pass filter for cortical processing^12,35,69^. Thus, in addition to enabling cortical circuits to adjust to recent history, this form of adaptation may also shape temporal integration by limiting the rate at which cortical circuits can follow rapidly fluctuating visual inputs, setting the threshold for flicker-fusion^70,71^.

Moving forward, we can begin to connect the diversity of perceptual effects of adaptation to the diversity of biological mechanisms that affect activity over time. Perceptual effects of adaptation can vary depending on duration even in response to a visual stimulus with the same spatial features. Our data indicate that this could arise through complementary mechanisms that ebb and flow on different time scales within the same neurons. This is consistent with studies that have identified multiple timescales of adaptation within single neurons that vary by orders of magnitude^9,10^. As a result, visual perception is shaped by concurrent dependencies on stimulus history that vary in their computational capacities. Another interpretation of these multiple forms of adaptation is as a series of mechanisms that work together to reduce activity in stages—if synaptic depression is not sufficient to reduce firing rates, cell-intrinsic hyperpolarization can reduce responses over longer periods of elevated excitation. Notably, this will come at the expense of stimulus specificity, but may be necessary to maintain cortical homeostasis. Indeed, prior studies using a greater number of stimulus presentations have identified both orientation specific and nonspecific components of adaptation^10,20^. Thus, future work will be important for understanding the nature of interactions between distinct mechanisms in individual neurons.

In summary, we have linked a well-studied synaptic mechanism to the *in vivo* phenomenon of adaptation at rapid timescales in V1. While distinct, synaptic and cell-intrinsic mechanisms need not be mutually exclusive and likely co-exist within single neurons^9,10,39^. Our findings provide long sought-after evidence for a synaptic depression mechanism at intracortical synapses that generates sensory adaptation and sparsens representations. Given the similarity of cortical structure and observed features of rapid adaptation across sensory areas, this mechanism could be applicable across many stimulus modalities^9,72–74^. Release probability is an inherent property at chemical synapses in the brain^11^; thus, molecular machinery and neuromodulators that affect Pr can regulate synaptic transmission at a given set of synapses over time (as we’ve studied here), but could also specialize the dynamics of responses across different brain areas, or even different species. Therefore, studies of short-term plasticity at synapses further along the visual hierarchy^20^, in different behavioral contexts^75^, sensory areas^9^, or species, could all generate insights into how fundamental attributes of synapses shape the neural code.

## Acknowledgements

We thank Gloria Kim for surgical assistance, TJ Wagner and Wenjuan Kong for husbandry assistance, Drs. Debra Silver and LJ Pilaz for equipment and instruction for performing *in utero* electroporation. We thank Drs. Celine Cammarata, Court Hull and Nicholas Priebe for comments on the manuscript, and members of the Hull and Glickfeld labs for insight throughout the project. This work was supported by grants from the National Eye Institute (R01-EY031328 to L.L.G. and F31-EY031941 to J.Y.L.).

## Author Contributions

Conceptualization: J.Y.L. and L.L.G.; Formal Analysis: J.Y.L.; Investigation: J.Y.L.; Data Curation: J.Y.L.; Writing-Original Draft: J.Y.L.; Writing-Review and Editing: J.Y.L. and L.L.G; Visualization: J.Y.L.; Supervision: L.L.G.; Funding Acquisition: J.Y.L. and L.L.G.

## STAR Methods

### Key Resources Table

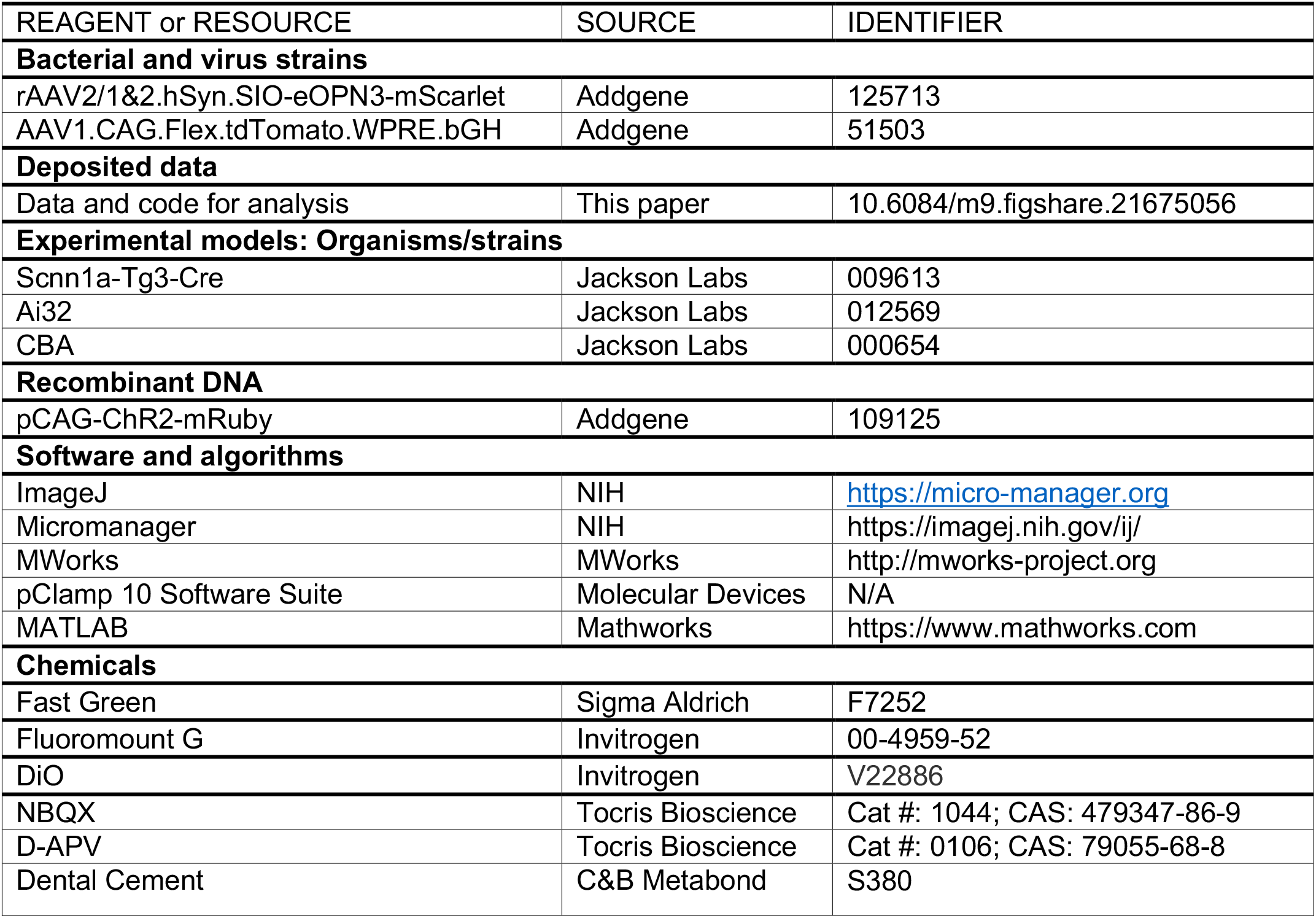

| REAGENT or RESOURCE | SOURCE | IDENTIFIER |
| --- | --- | --- |
| **Bacterial and virus strains** |  |  |
| rAAV2/1&2.hSyn.SIO-eOPN3-mScarlet | Addgene | 125713 |
| AAV1.CAG.Flex.tdTomato.WPRE.bGH | Addgene | 51503 |
| **Deposited data** |  |  |
| Data and code for analysis | This paper | 10.6084/m9.figshare.21675056 |
| **Experimental models: Organisms/strains** |  |  |
| Scnn1a-Tg3-Cre | Jackson Labs | 009613 |
| Ai32 | Jackson Labs | 012569 |
| CBA | Jackson Labs | 000654 |
| **Recombinant DNA** |  |  |
| pCAG-ChR2-mRuby | Addgene | 109125 |
| **Software and algorithms** |  |  |
| ImageJ | NIH | https://micro-manager.org |
| Micromanager | NIH | https://imagej.nih.gov/ij/ |
| MWorks | MWorks | http://mworks-project.org |
| pClamp 10 Software Suite | Molecular Devices | N/A |
| MATLAB | Mathworks | https://www.mathworks.com |
| **Chemicals** |  |  |
| Fast Green | Sigma Aldrich | F7252 |
| Fluoromount G | Invitrogen | 00-4959-52 |
| DiO | Invitrogen | V22886 |
| NBQX | Tocris Bioscience | Cat #: 1044; CAS: 479347-86-9 |
| D-APV | Tocris Bioscience | Cat #: 0106; CAS: 79055-68-8 |
| Dental Cement | C&B Metabond | S380 |

## RESOURCE AVAILABILITY

### Lead contact

Further information and requests for resources and reagents should be directed to Lindsey Glickfeld.

### Materials availability

No new reagents were generated as a result of this study.

### Data and code availability

- All electrophysiology data included in the manuscript figures is available on Figshare. A link is provided in the *Key resources table*.
- All original code needed to generate the manuscript figures is available on Figshare. A link is provided in the *Key resources table*.
- Any additional information required to reanalyze the data reported in this paper is available from the lead contact upon request.

## EXPERIMENTAL MODEL AND SUBJECT DETAILS

### Animals

All procedures conformed to standards set forth by the National Institutes of Health Guide for the Care and Use of Laboratory Animals, and were approved by the Duke University’s Animal Care and Use Committee. Mice were housed on a normal 12:12 light-dark cycle. Data in this study were collected from 74 mice (35 female). For experiments involving selective expression in layer 4 V1 neurons, we used Cre-positive offspring from Scnn1a-Tg3-Cre mice (Jackson Labs #009613) crossed with either Ai32 (Jackson Labs #012569, n = 15), or CBA (Jackson Labs, #000654, n = 18). We also used offspring from Scnn1a-Tg3-Cre and CBA mice for *in utero* electroporation (n = 11) but did not select for Cre expression. All other experiments did not require cell-type specific expression; thus, mice were a mix of genotypes (n = 32). Transgenic mice were heterozygous and bred on a C57/B6J background (Jackson Labs #000664) with up to 50% CBA/CaJ (Jackson Labs #000654). *In vivo* electrophysiology experiments used mice 6-22 weeks old and *in vitro* electrophysiology experiments used mice 4-12 weeks old. At the time of viral injection, mice were at least 4 weeks old.

## METHOD DETAILS

### Surgical Procedures

#### Intracranial viral injections

Burrhole injections of viral constructs [rAAV2/1&2.hSyn.SIO-eOPN3-mScarlet (Addgene 125713 diluted to 6 × 10^12^ viral genomes/mL) or AAV1.CAG.Flex.tdTomato.WPRE.bGH (Addgene 51503; diluted to 3 × 10^12^ viral genomes/mL)] were used to selectively express opsins and control fluorophores in layer 4. Mice were anesthetized with isoflurane and positioned in a stereotax (Kopf Instruments). Meloxicam (5 mg/kg) was administered subcutaneously and bupivacaine (5 mg/kg) was administered locally prior to incision. After the skull was exposed, a small hole was drilled -2.6 mm lateral from lambda and directly anterior to the lambdoid suture targeting the posterior and medial aspect of the primary visual cortex (V1). Injection micropipettes were pulled from glass capillary tubes (1B100F-4, World Precision Instruments) and backfilled with virus and then mineral oil and mounted on a Hamilton syringe. The pipette was lowered into the brain and pressure injected at two depths using an UltraMicroPump (World Precisions Instruments; 2 × 100 nL; -350 μm and -450 μm from the surface). We waited between 4.5-7 weeks for viral expression for both *in vitro* and *in vivo* electrophysiology and confirmed expression *post hoc*.

#### In utero electroporation

Embryos from timed-pregnant CBA female mice (E15.5-16.5) mated to Scnn1a-Tg3-Cre males were used to obtain expression in layer 2/3. Meloxicam was administered pre-operatively (1 mg/mL, 5 mg/kg; subcutaneous). Animals were maintained under anesthesia (2.5% isoflurane), the abdomen was cleaned with ethanol and then swabbed with iodine. An incision was made in the skin and then in the abdominal wall, then covered in a drape made with sterile surgical gauze. Uterine horns were carefully removed and kept moist with warm PBS throughout the surgery. Embryos were injected with plasmid mixture (1.5 μg/uL pCAG-ChR2-mRuby-ST in 0.5% Fast Green in UltraPure water, Addgene 109125) in the left ventricle using a glass micropipette pulled to a 70 μm beveled tip. After injection, a series of voltage steps (five voltage pulses of 50 V at 1 Hz with each pulse lasting 50 ms) was applied to each embryo using 5 mm round tweezertrodes (BTX, BTX ECM 830 ElectroSquarePorator). Paddles were oriented to target V1. Embryos were gently returned to the abdomen in the same side that they were removed from. The abdominal wall was sutured before applying bupivicane (5 mg/kg) and then suturing the skin. Animals were allowed to recover on a heating pad until mobile. Strength and location of expression was screened with trans-cranial fluorescence of mRuby following headpost implantation.

#### Headpost implantation

Mice were anesthetized with a mixture of ketamine/xylazine (ketamine: 50 mg/kg, xylazine: 5 mg/kg; intraperitoneal) and isoflurane (1.2–2% in 100% O_2_). Meloxicam was administered pre-operatively (1 mg/mL, 5 mg/kg; subcutaneous). Using aseptic technique, a custom-made titanium headpost was secured over V1 using clear dental cement (C&B Metabond, Parkell). Buprenex (0.05 mg/kg) and cefazolin (50 mg/kg) were administered post-operatively. Animals were allowed to recover for at least 1 week prior to experiments.

### Visual and optogenetic stimulus presentation

Visual stimuli were presented on a 144-Hz (Asus) LCD monitor, calibrated with an i1 Display Pro (X-rite). The monitor was positioned 21 cm from the contralateral eye. Visual stimuli were controlled with MWorks (http://mworks-project.org). Circular gabor patches containing sine-wave gratings (30º diameter; 0.1 cycles per degree; 80% contrast) alternated with periods of uniform mean luminance (60 cd/m^2^). Timing of visual stimulus onset was measured for aligning neural data via a photodiode that directly measured output from the LCD. All baseline and test stimuli were presented for 0.1 s, with inter-stimulus intervals (ISIs) ranging from 0.25 s to 4 s and inter-trial interval of 8 s to allow for adequate recovery.

#### ChR2 activation

Control and ChR2 activation trials were randomly interleaved. Control trials consisted of two vertically oriented static gratings separated by a 0.25, 1, or 4 s ISI. ChR2 activation trials consisted of a sine-wave laser pulse (0.5 s, 20 Hz, 450 nm, Optoengine) followed by a grating (0.25, 1, or 4 s ISI). In a subset of experiments, two static gratings (0.25 s ISI) were presented following ChR2 activation to measure the effect on visual adaptation. The effect of serial ChR2 activation was also tested using brief (0.1 s) square-wave pulses (**Figure S6A-C**).

#### Stimulus specificity of adaptation

Two protocols were used to test the stimulus specificity of adaptation. 1) Five repeated presentations of a static grating (baseline and test 1-4; 0.25 s ISI) followed by a presentation of the orthogonal orientation (test 5). On randomly interleaved trials, the repeated and orthogonal orientation were switched to obtain the baseline amplitude of both orientations. 2) Two oriented gratings were presented with an ISI of 0.25 s. The test stimulus was the same across trials while the baseline stimulus was varied from 0 to 90 degrees from the test in 22.5 degree increments.

#### Orientation tuning

Drifting gratings (2 Hz) moving in 16 directions (22.5 degree increments) were presented for 1 s with an 8 s inter-trial interval to measure the orientation tuning of neurons.

#### eOPN3 activation

All trials consisted of two vertically oriented baseline and test stimuli separated by 0.25 s ISI. After 20 control trials, eOPN3 was activated with a square-wave laser pulse (10 s, 10 Hz, 530 nm, Optoengine). We then tested the effect of eOPN3 over 20 trials with top-up activation (0.5 s, 10 Hz) preceding visual stimulation on each trial. Recovery was measured during a subsequent 90-100 trials. Each experiment contained 1-3 repeats of eOPN3 activation blocks.

#### Contrast dependence of adaptation

Two vertically oriented gratings were presented with an ISI of 0.25 s. The test stimulus was the same (80% contrast) across trials while the baseline stimulus was either 40 or 80% contrast.

### Experimental Procedures

#### In vivo retinotopic mapping

For all *in vivo* electrophysiological recordings, V1 boundaries were first identified with retinotopic mapping with intrinsic signal imaging through the skull. The skull was illuminated with orange light (590 nm LED, Thorlabs), and unfiltered emitted light was collected using a CCD camera (Rolera EMC-2, Q Imaging) at 2 Hz through a 5x air immersion objective (0.14 numerical aperture (NA), Mitutoyo), using Micromanager acquisition software^76^. Drifting gratings (80% contrast, 2 Hz, 0.1 cpd) were presented for 2 s at 3 positions with a 4 s interstimulus interval. Collected images were analyzed in ImageJ to measure changes in reflectance at each position (dR/R; with R being the average of all frames) to identify V1.

#### Preparation for in vivo electrophysiology

Animals were habituated to head-fixation for 1-3 days prior to surgery. On the day of recording, animals were anesthetized with isoflurane and a small craniotomy (< 1 mm diameter) was made over a V1 location identified by intrinsic signal imaging. For extracellular recordings, a gold ground pin was inserted in an anterior portion (outside of visual areas) within the headpost and secured with dental cement. Damage to superficial cortex was minimized by drilling in brief bouts (< 1 s) and alternating drilling and cooling with chilled glucose-free HEPES-based artificial cerebral spinal fluid (ACSF; in mM: 141 NaCl, 2.5 KCl, 10 HEPES, 2 CaCl_2_ 1.3 MgCl_2_). A slit was made in the dura with a syringe and the craniotomy was kept covered with ACSF for the remainder of the experiment. Animals were allowed to recover on the running wheel for at least 45 minutes before recording. In a subset of experiments, recording was performed the day after the craniotomy or animals were used for up to 2 consecutive recording days. In these cases, the craniotomy was protected overnight with Dura-Gel (Cambridge NeuroTech) and dental cement, which were removed and replaced with ACSF prior to recording.

#### In vivo whole-cell recordings

Whole-cell recordings were performed using blind patch technique. A silver chloride ground pellet was placed in the recording well outside of the brain. Recording ACSF was wicked away from the craniotomy and a 3-5 MOhm glass micropipette was lowered until the pipette tip touched the brain (confirmed by appearance of a square pulse on the membrane test); this position was zeroed and the well was refilled with recording ACSF. All recordings were documented relative to this depth. The pipette was lowered to ∼100 μm depth and then stepped in 1-2 μm increments until an increase in resistance was observed and pressure was released to form a GΩ seal. Cells recorded at 180-350 μm depths were considered to be within L2/3. For current clamp recordings, internal solution contained (in mM): 142 K-gluconate, 3 KCl, 10 HEPES, 0.5 EGTA, 5 phosphocreatine-di(tris), 5 phosphocreatine-Na_2_, 3 Mg-ATP, 0.5 GTP; for voltage clamp recordings, internal solution contained (in mM): 125 Cs-methanesulfonate, 5 TEA-Cl, 10 HEPES, 0.5 EGTA, 4 Mg-ATP, 0.3 Na_3_GTP, 8 phosphocreatine-di(tris), 3 NaCl. For voltage clamp recordings, EPSCs were recorded at -70 mV and IPSCs were recorded at +10 mV based on previous literature and our own calibration with ChR2 activation of interneurons *in vivo* (**Figure S2A**). Series resistance was monitored using -5 mV steps preceding each stimulus; recordings that reached >35 MΩ resistance or >20% change from baseline were discarded. The order of recording EPSCs and IPSCs was varied across experiments, and there was no relationship between the series resistance and the normalized current for either holding potential (**Figure S2B-C**; EPSCs p = 0.21, IPSCs p = 0.57).

In a subset of recordings, a low resistance pipette (1 MΩ) was filled with 3 M NaCl and lowered ∼200 μm to measure local field potential and determine optimal stimulus position. Otherwise, optimal stimulus position was determined separately for azimuth and elevation by observing spikes or EPSCs in response to a flashing white bar (0.1 s on, 1 s off, 5 degree width). Following optimization of stimulus position, spikes and EPSCs were analyzed online to determine the preferred stimulus orientation.

#### In vivo extracellular recordings

Extracellular recordings were performed with a 32-site acute probe (A1×32-Poly2–5mm-50s-177-A32, NeuroNexus or H4, Cambridge NeuroTech). Probes were connected through an A32-OM32 adapter to a Cereplex Mu digital headstage (Blackrock Microsystems). Signals were digitized at 30 kHz and recorded by a Cerebus multichannel data acquisition system (Blackrock Microsystems). Probes were slowly lowered into the brain until all sites were inserted and allowed to stabilize for 40–50 min before recording. For experiments involving localized viral expression, the probe was painted with DiO (Thermo Fisher) to confirm with *post hoc* histology that the electrode tract was within the expression region.

For optogenetic experiments, we used either a 450 nm or 532 nm laser (Optoengine) to activate ChR2-expressing neurons or inhibit L4 terminals with eOPN3, respectively. Lasers were coupled to an optic shutter and patch cable terminating in an optic fiber. For L2/3 ChR2 activation and eOPN3 inhibition experiments, probes had attached optic fibers (200 μm core, 0.22 NA) that terminated 100 μm above the surface of the brain. For L4 ChR2 stimulation, a tapered lambda fiber (100 μm core with 0.9 mm taper, 0.22 NA, Optogenix) was inserted in the brain aligned to the tip of the probe for enhanced light transmission deeper in the brain. Laser power was calibrated to deliver 1 mW power at fiber tip for ChR2 activation and 1.2 mW power at the fiber tip for eOPN3 inhibition.

On a subset of recordings, putative ChR2-expressing units were identified by blocking excitatory transmission with a mix of AMPAR and NMDAR blockers (3 mM NBQX and 6 mM APV, respectively) diluted in 100 μL of recording ACSF^77^. At the end of the recording, ACSF was wicked away from the recording well and the drug mixture was dripped onto the craniotomy. After at least 20 minutes (up to 45 minutes, based on visually-evoked responses at the deepest electrode sites) ∼50 pulses of 450 nm laser (10 ms, 0.1 Hz) were presented to activate ChR2-expressing cells.

#### In vitro slice preparation

Mice were deeply anesthetized with isoflurane, the brain was removed and then transferred to oxygenated (95% O_2_ and 5% CO_2_), ice-cold artificial cerebrospinal fluid (ACSF, in mM: NaCl 126, KCl 2.5, NaHCO_3_ 26, NaH_2_PO_4_ 1.25, glucose 20, CaCl_2_ 2, MgCl_2_ 1.3). Coronal brain slices (300 μm thickness) were prepared using a vibrating microtome (VT1200S, Leica) and transferred to a holding solution (at 34º C) for 12 minutes, and then transferred to storage solution for 30 min before being brought to room temperature. The holding solution contained (in mM): 92 NaCl, 2.5 KCl, 1.25 NaH_2_PO_4_, 30 NaHCO_3_, 20 HEPES, 25 glucose, 2 thiourea, 5 Na-ascorbate, 3 Na-pyruvate, 2 CaCl_2_, 2 MgSO_4_. The storage solution contained (in mM): 93 NMDG, 2.5 KCl, 1.2 NaH_2_PO_4_, 30 NaHCO_3_, 20 HEPES, 25 glucose, 2 thiourea, 5 Na-ascorbate, 3 Na-pyruvate, 0.5 CaCl_2_, 10 MgSO_4_. Micropipettes pulled from borosilicate glass (1B150F-4, World Precision Instruments) were filled with internal solution containing (in mM): 142 K-gluconate, 3 KCl, 10 HEPES, 0.5 EGTA, 5 phosphocreatine-tris, 5 phosphocreatine-Na2, 3 Mg-ATP, 0.5 GTP. Recording pipettes had resistances of 3-5 MΩ.

#### In vitro slice recordings

Recordings occurred between 1.5 and 5 hours after the animal was sacrificed. Brain slices were transferred to a recording chamber and maintained at 34º C in oxygenated ACSF perfused at 2 mL/min. Electrophysiological recordings were restricted to layer 2/3 and V1 was identified either by reference atlas alignment or visualization of fluorescence expression at the viral injection site. Neural signals were recorded using a MultiClamp 700B and digitized with a Digidata 1550 (Axon Instruments) with a 20 kHz sample rate. Data acquisition and stimulus presentation was controlled using the Clampex software package (pClamp 10.5, Axon Instruments).

In current-clamp recordings, a constant positive current was injected to maintain membrane potential near resting membrane potential measured *in vivo*. To test effects of depolarization on membrane potential, positive current was injected for a duration that varied between 0.1 and 5 s. Current level was calibrated with 0.1 s current injections to elicit a similar firing rate (∼30 Hz) across cells, but generally ranged between 400-600 pA.

In voltage-clamp recordings, series resistance was monitored using -5 mV steps preceding each trial. At least 10 sweeps were collected for each recording condition. Only cells that had < 20 MΩ series resistance, < 20% series resistance change, and stable holding current (<100 pA baseline variation) were included for analysis. EPSCs were evoked by either electrical stimulation with a steel monopolar electrode placed in L4 or L2/3 (100 μs pulse) or optical activation of ChR2 over cell bodies in L4 (light power 0.5-1.5 mW/mm^2^, 470 nm LED, 100 ms square pulse). For optical activation, light pulses from a 4-color LED controller (ThorLabs) were coupled to the epifluorescence path (Olympus BX-RFA) and projected through a 40x water immersion lens (Olympus, 0.8 NA). To minimize polysynaptic activation, electrical and optical stimulation intensities were calibrated to elicit EPSCs with ∼100-250 pA amplitude and short latency (< 5 ms).

Since electrical stimulation activates axons non-selectively, in a separate set of experiments in Scnn1a-Cre x Ai32 animals we compared EPSCs in L2/3 neurons in response to electrical and optogenetic activation in L4. After patching a L2/3 neuron, a small spot (50 μm, 100 μs) of 470 nm light was used to search for an area in L4 that elicited short-latency, monosynaptic responses. The stimulation electrode was placed in the center of this spot, presumably near a L4 neuron synapsing onto the L2/3 neuron being recorded. EPSCs recorded in L2/3 neurons displayed the same depression for electrical and optogenetic stimulation, indicating that L4 electrical stimulation is sufficient to reveal the dynamics of L4-L2/3 synapses (**Figure S4**).

eOPN3 in L4 terminals was activated by illuminating a small area (100 μm diameter) around the recorded neuron with green light (0.8 mW/mm^2^, 530 nm LED) for 10 s, followed by a 0.5 s top-up preceding each trial.

#### Post hoc histology

After recording in virally injected or electroporated animals, brains were imaged to confirm viral expression in the recorded area. For *in vitro* recordings, slices were incubated 12-16 hours in 4% paraformaldehyde (PFA) in PBS, washed 3x with PBS and mounted. For *in vivo* recordings, the probe tract was visualized with DiI or DiO painted on the probe prior to insertion (Invitrogen V22889). After recording, animals were anesthetized with an overdose of ketamine/xylazine and perfused with PBS followed by 4% PFA in PBS. Brains were dissected and incubated in 4% PFA overnight, rinsed 3x with PBS, then sliced in 100 μm sections and mounted on glass slides. Slides were mounted with Fluoromount G with DAPI (Invitrogen) and imaged using a Zeiss inverted microscope (Zeiss Axiovert).

## QUANTIFICATION AND STATISTICAL ANALYSIS

All analyses were performed in custom code written in either MATLAB or Python. All data are presented as mean ± SEM. N values refer to number of cells or units isolated. Sample sizes were not predetermined but are comparable to published literature for each type of experiment. For all experiments adaptation is quantified as the normalized response:

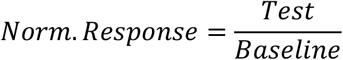

Where the *Baseline* response is the response to the first visual stimulus (or electrical or optical stimulus *in vitro)* on a trial and the *Test* is the response to the same stimulus following a visual, electrical, or optical adapter.

### Analysis of *in vivo* whole-cell recordings

#### Current clamp recordings

Raw membrane potential was separated into firing rate and subthreshold membrane potential. Firing rate was obtained by setting a voltage threshold on a cell-by-cell basis for detecting spikes. Subthreshold membrane potential was obtained by using a median filter to clip spikes. For each ISI, pre-stimulus mean membrane potential and variance were measured from subthreshold membrane potential in a 0.1 s window prior to stimulus onset. Spike threshold was measured from spikes detected in a 0.4 s window around stimulus onset (0.1 s before and 0.3 s after stimulus onset). Spike threshold was calculated by averaging over the membrane potential at the time of the peak of the second derivative for all spikes within this time window. PSP amplitude was measured in a 20 ms window around the peak of the trial-averaged response during the stimulus-evoked response window (0-0.25 s after stimulus onset), relative to the baseline window (0.1 s before stimulus onset).

#### Voltage clamp recordings

EPSC or IPSC stimulus-evoked amplitude was quantified by averaging current values within a 20 ms window around the peak of the response in the stimulus-evoked response window (0-0.25 s after stimulus onset). Mean and standard deviation of the holding current was quantified in a 0.1 s window prior to stimulus onset. Recovery time constants were fit for EPSCs and IPSCs using a single exponential from the normalized current amplitude averaged across cells. For all stimulus specificity experiments, current amplitudes were normalized to the baseline stimulus of the same orientation. Adaptation tuning width was measured by fitting the normalized responses with a von Mises function.

### Analysis of *in vitro* whole-cell recordings

Amplitudes of EPSCs in response to electrical stimulation or 0.01 s ChR2 activation were calculated as the mean of the trial-averaged response in a 2 ms window around the peak of the response. Amplitudes of EPSCs in response to 0.1 s ChR2 activation were calculated in a 20 ms window around the peak of the response. Recovery of optogenetically evoked EPSCs from adaptation with 0.1 s ChR2 activation was fit with a single exponential.

### Analysis of extracellular recordings

#### Spike sorting

Single units were isolated with KiloSort 2.5 (https://github.com/MouseLand/Kilosort) using refractory period violations and steepness of the autocorrelogram as criteria for isolation. We then manually curated these units in Phy (https://github.com/cortex-lab/phy) such that only units that were detected throughout the entire recording are included for subsequent analysis. Depth of the unit was assigned based on their waveforms’ center-of-mass across sites. Fast-spiking (FS) and regular-spiking (RS) units were separated within recordings according to peak-to-trough time of the maximum amplitude waveform across all contact sites (**Figure S3**).

#### Layer identification

To functionally identify cortical layers, we used the local field potential (LFP) obtained from filtering the raw data (downsampled to 10 kHz) from 1 to 200 Hz. The trial-averaged, stimulus-evoked LFP during a 1-second drifting grating presentation was converted to a current source density (CSD) plot by taking the discrete second derivative across the electrode sites and interpolated. Layer bounds were assigned relative to an initial sink in layer 4, followed by a sink in layer 2/3 and a sustained sink in layer 5 (**Figure S5**).

To confirm layer identification, in a subset of experiments ChR2-activated units were identified in the presence of excitatory synaptic blockers to identify ChR2-expressing units. Each unit’s distance in depth from the L23-L4 boundary was measured to compare depth of L4 versus L2/3 ChR2-expressing neurons across the two experiment types (**Figure S5**).

#### Data inclusion and analysis

For all recordings, only cells that were visually responsive, according to a paired t-test in a 0.15 s window before and after stimulus onset, were included. In ChR2 activation experiments, “laser active” units were defined as units significantly driven by ChR2 activation in this same time window. For eOPN experiments, inhibited and facilitated units were defined as having >20% decrease or increase, respectively, in visually-evoked responses during eOPN3 activation compared to control trials. Categorization of units with significant increase or decrease defined by paired t-test yielded similar results. Neurons that were classified as inhibited in L2/3 and L4 were monitored for recovery of visually-evoked responses following eOPN3 activation and the recovery time constant was fit with a single exponential from the start of eOPN3 induction.

PSTHs were generated by binning spiking activity in 0.01 s windows across all trials of each type, aligned to stimulus onset. Each stimulus condition contained at least 20 repeats. Maximum firing rate was measured as the average firing rate in a 20 ms window around the peak of the PSTH. For plots visualizing average stimulus-evoked responses across units, firing rates were z-scored prior to averaging. Orientation tuning was measured using the mean firing rate in a 20 ms window around the peak PSTH for each stimulus direction, collapsed by orientation and fit with a von Mises function.

## Supplemental Figures

**Figure S1.**
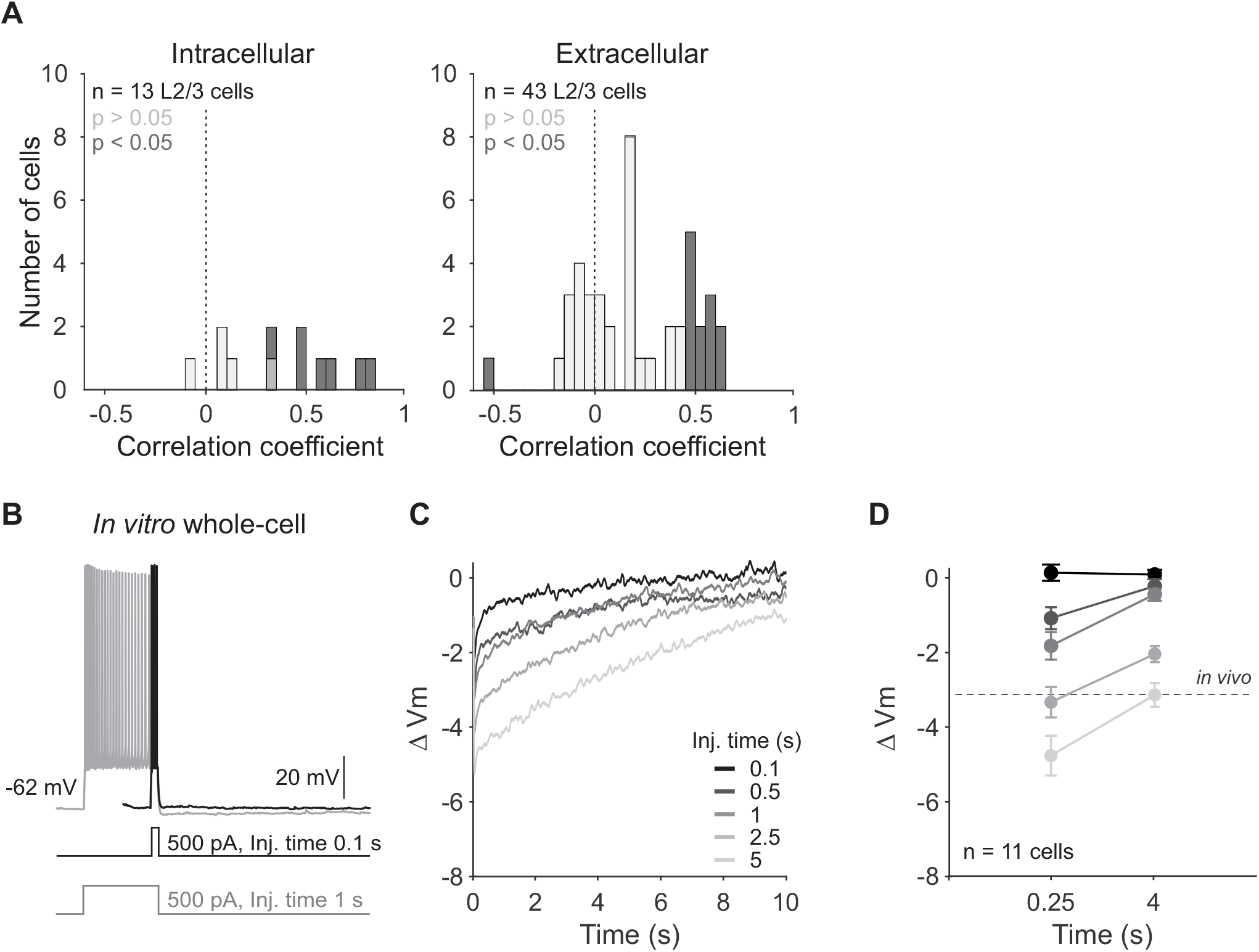
Rapid adaptation does not induce cell-intrinsic fatigue, related to Figure 1. **A**. Histogram of single trial correlation of spikes elicited in response to baseline and test stimuli for 0.25 ISI condition from intracellular (left; n = 13 cells) and extracellular (right; n = 43 units) L2/3 *in vivo* recordings. Dark gray bars indicate significant correlations. **B**. Current clamp recording in example L2/3 pyramidal cell in response to current injections of two durations (black = 0.1 s, gray = 1 s). **C**. Change in membrane potential (Vm) following offset of increasing current injection durations in the example cell in **B. D**. Average change in membrane potential after current injection offset at recovery times when spike output is suppressed (0.25 s) or recovered (4 s) *in vivo* for increasing current durations. Dashed line is average change in stimulus-evoked membrane potential *in vivo*. Error bar is SEM across cells

**Figure S2.**
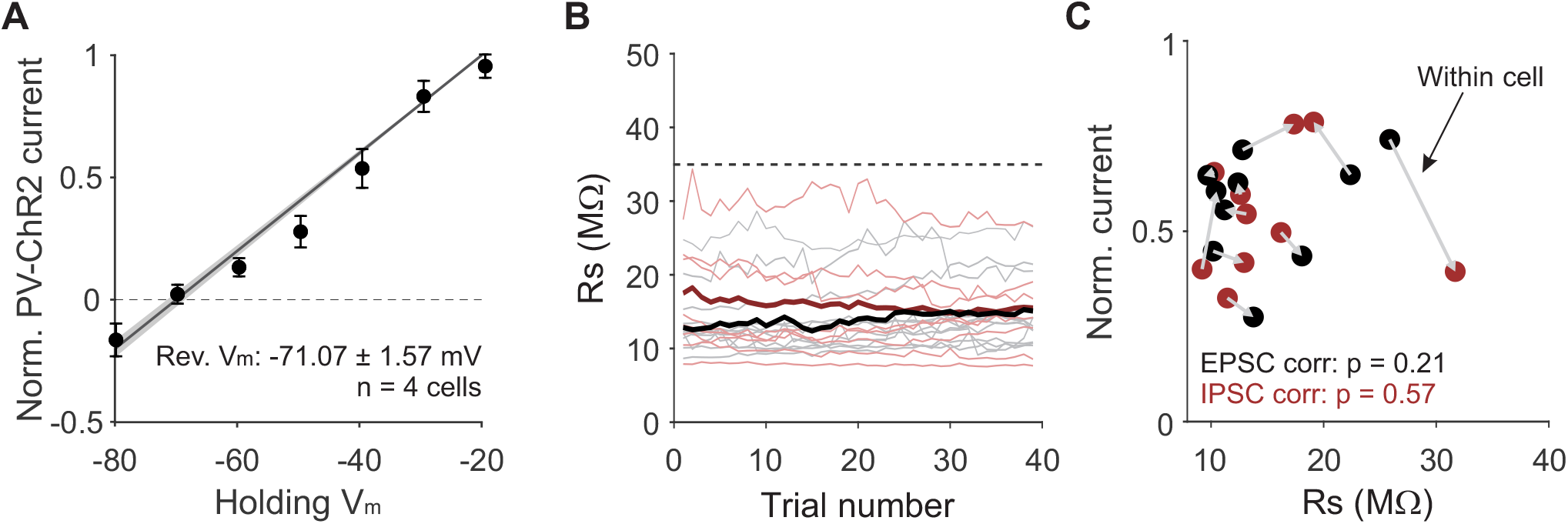
Whole-cell voltage clamp recording of EPSCs and IPSCs *in vivo*, related to Figure 2. **A**. Reversal potential of currents evoked with optogenetic activation of parvalbumin-expressing (PV) interneurons expressing Channelrhodopsin-2 (ChR2) to calibrate reversal potential for inhibitory currents in vivo (PV-Cre mice injected with AAV2/1.hSyn.ChR2-YFP; n = 4 cells). **B**. Series resistance (Rs) during recording of EPSCs (black) and IPSCs (red). Thick lines are average across cells. Dashed line is cutoff used for series resistance inclusion criteria. **C**. Normalized current (baseline/test; 0.25 s ISI) as a function of series resistance for all recordings. Grey arrows connect currents recorded within the same cell and direction reflects the order of recording. P-value is significance of Pearson correlation.

**Figure S3.**
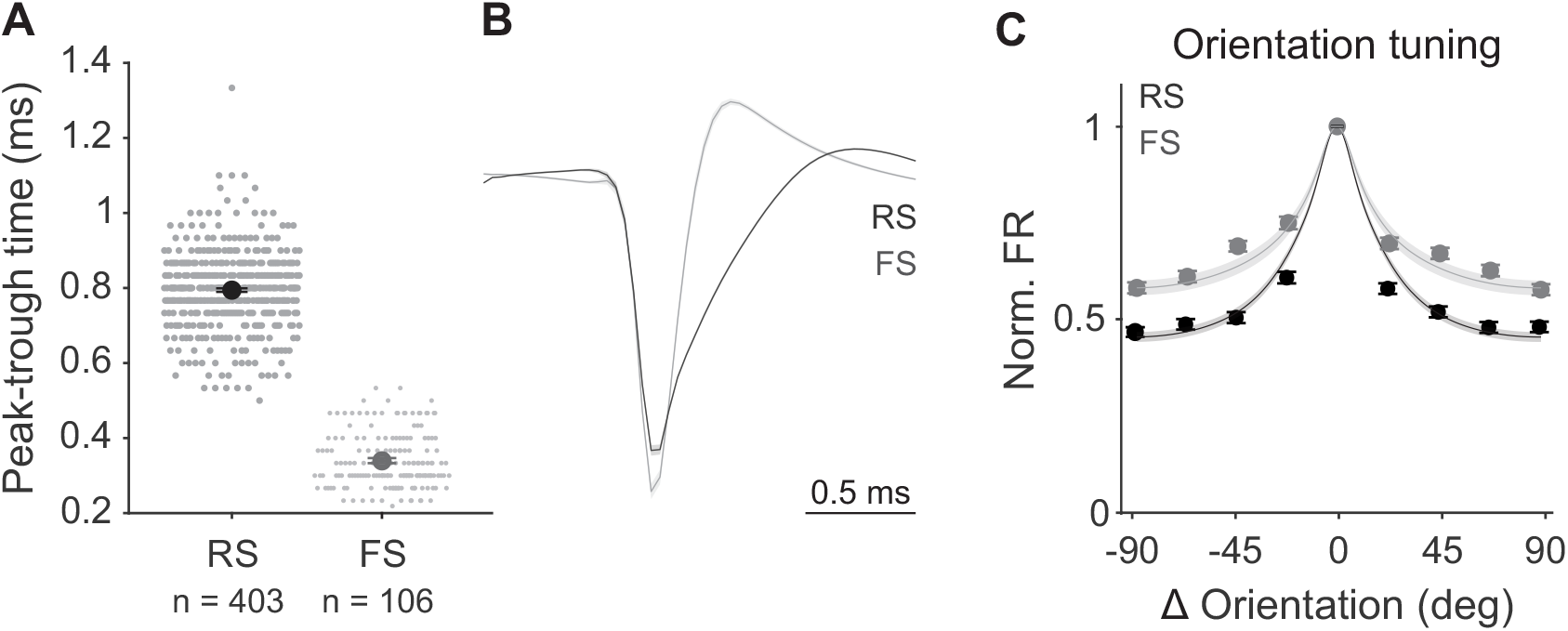
Separation of FS and RS units and comparison of orientation tuning, related to Figures 2 and 3. **A**. Peak-trough time of spike waveforms from units classified as regular spiking (RS, black) or fast spiking (FS, grey). **B**. Average spike waveforms from the units in **A**. Shaded error is SEM across units. **C**. Average orientation tuning curves aligned to preferred orientation for each unit. Points are averaged normalized response across units. Curves are averages of the von Mises fit for individual units.

**Figure S4.**
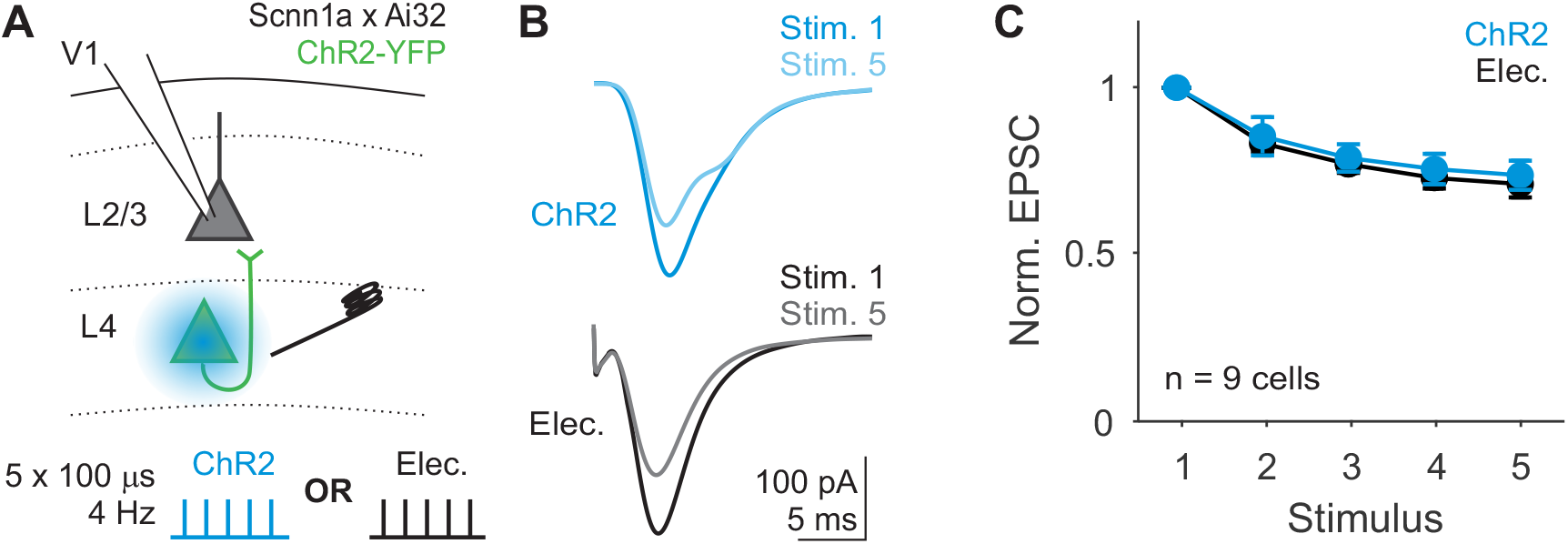
EPSCs in L2/3 measured with ChR2 and electrical stimulation, related to Figure 4. **A**. Schematic of recording EPSCs from a L2/3 pyramidal cell while stimulating L4 neurons optogenetically or electrically on alternating trials. **B**. Average EPSCs from an example cell in response to optogenetic (blue) or electrical (black) stimulation of L4 for the first (dark) and last (light) stimulus in the train. **C**. Average EPSC amplitude normalized to first pulse within stimulation type. Error bar is SEM across cells. Two-way ANOVA, p = 0.51 for effect of stimulation type.

**Figure S5.**
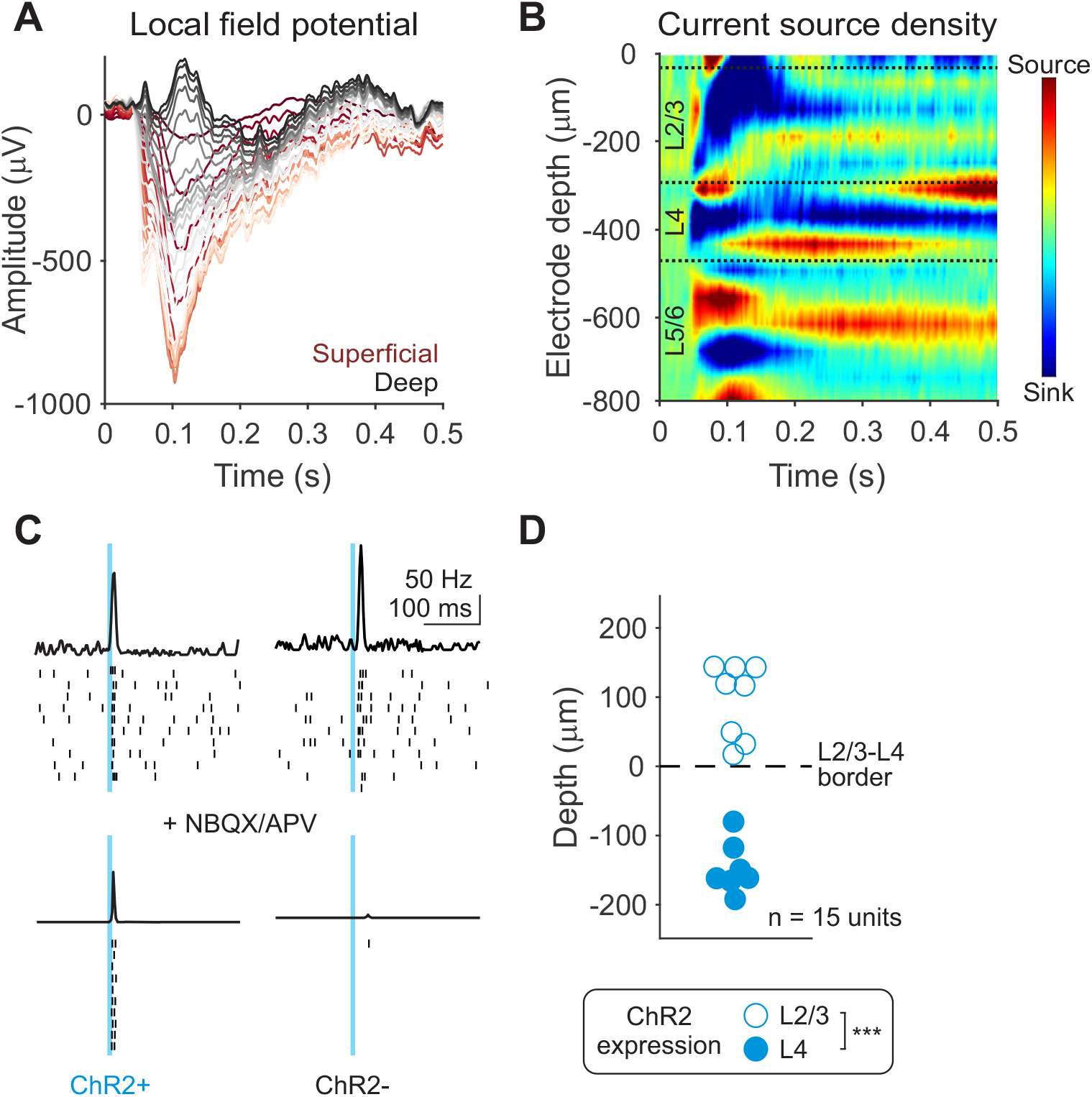
Identification of layer boundaries for classifying units as L2/3 or L4, related to Figures 5, 6 and 8. **A**. Local field potential (LFP) measured across cortical depths during a drifting grating stimulus from an example recording. Traces are colored according to contact site from superficial (red) to deep (black). **B**. Current source density (CSD) calculated using the LFP in **A**. Dashed lines indicate layer boundaries assigned based on this map. L4 was assigned by identifying an early onset sink and L2/3 was identified as the later onset sink above it. **C**. Example units identified as ChR2+ (left) or ChR2-(right). PSTH and spike rasters in response to blue laser pulses (10 ms) before (top) and after (bottom) pharmacological block of excitatory transmission (**STAR Methods**). **D**. Depth of ChR2-expressing units relative to L2/3-L4 boundary identified using the CSD. Marker fill indicates ChR2 expression layer (unfilled = L2/3, in utero electroporated mice; filled = L4, Scnn1a x Ai32 mice; depth of L2/3 vs L4 expression: p < 0.001, un-paired t-test).

**Figure S6.**
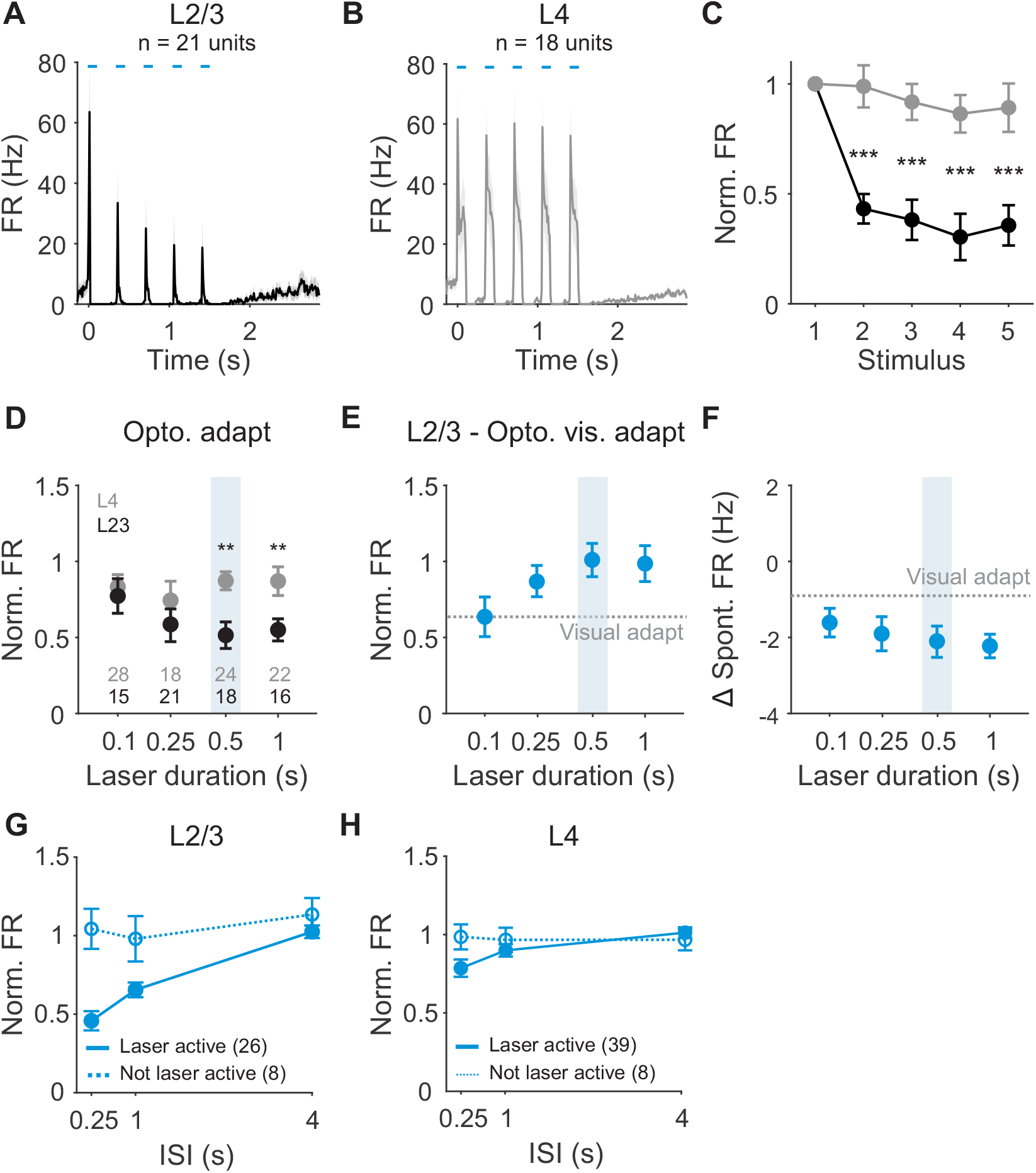
Effects of optogenetic activation of L4 neurons, related to Figure 5. **A**. Average PSTH of laser active L2/3 units during optogenetic activation of L4 neurons with 5, 0.1 s square pulses of blue light. **B**. Same as **A**, for L4 units. **C**. Peak firing rate for each stimulus pulse, normalized to the first pulse in the train for L2/3 (black) and L4 (grey) units. Error bar is SEM across units. L2/3 p < 0.001 for stimulus 2-5 vs 1, one-way ANOVA with post hoc Tukey test. **D**. Optogenetic adaptation measured in L2/3 and L4 units across different laser durations (L4 vs L2/3, 0.5 s laser duration: p = 0.006; 1 s laser duration: p = 0.009, unpaired t-tests). Shaded box indicates laser duration used for main figure experiments. **E**. Visual adaptation measured in L2/3 neurons following different durations of optogenetic activation of L4 neurons. Dashed line is visual adaptation in the absence of L4 stimulation. **F**. Change in spontaneous firing rate after different durations of L4 activation. Dashed line is spontaneous firing rate after visual adaptation. **G**. L2/3 optogenetic adaptation in units divided by units that were significantly modulated by L4 ChR2 stimulation (solid line) or not (dashed line). **H**. Same as **G**, for L4 units.

**Figure S7.**
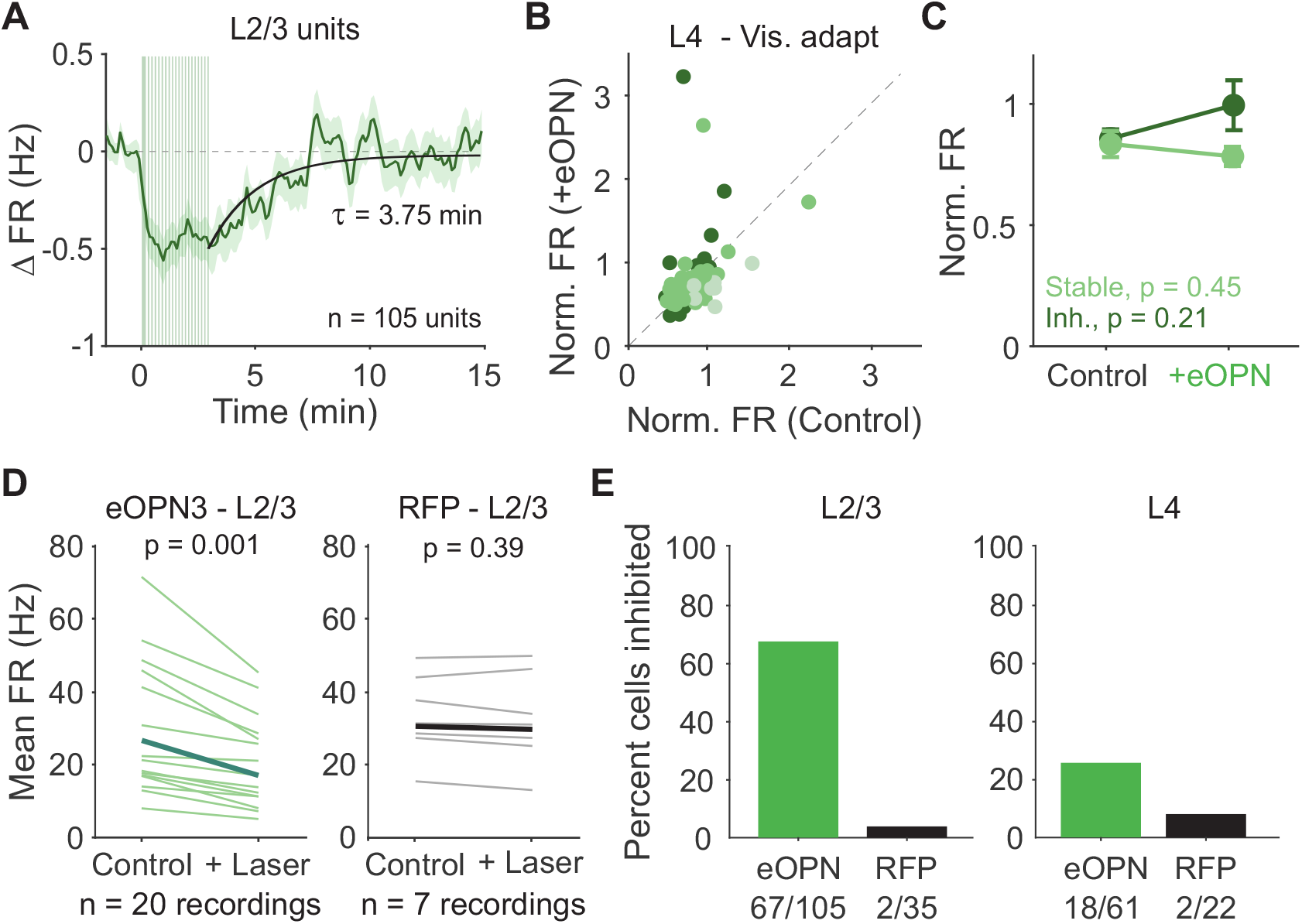
Green laser alone does not affect firing rates, related to Figure 8. **A**. Average time course of stimulus-evoked, z-scored firing rate aligned to eOPN3 activation for all units recorded in L2/3 (n = 105). Green vertical lines indicate eOPN3 activation trials. Black curve is fit to the recovery from eOPN3 activation. Shaded error is SEM across units. **B**. Comparison of normalized response (test/baseline) in control and eOPN3 activation trials, for all L4 units colored by categorization in **Figure 8E** (dark green = inhibited, medium green = stable, light green = facilitated). **C**. Average normalized response for inhibited (dark green) and stable (light green) units in L4. Error bar is SEM across units. **D**. Average visually-evoked firing rate of L2/3 neurons during control and laser stimulation trials in eOPN3 (left, green) or RFP control (right, black) recordings. Individual lines are average response of all L2/3 neurons in each session, thick line is mean across sessions (eOPN: paired t-test, p < 0.001; RFP: paired t-test p = 0.39). **E**. Left: Fraction of L2/3 units classified as inhibited from recordings with eOPN3 (green) or RFP control (black) in L4 neurons. Right: Same as left, for L4 units.

**Figure S8.**
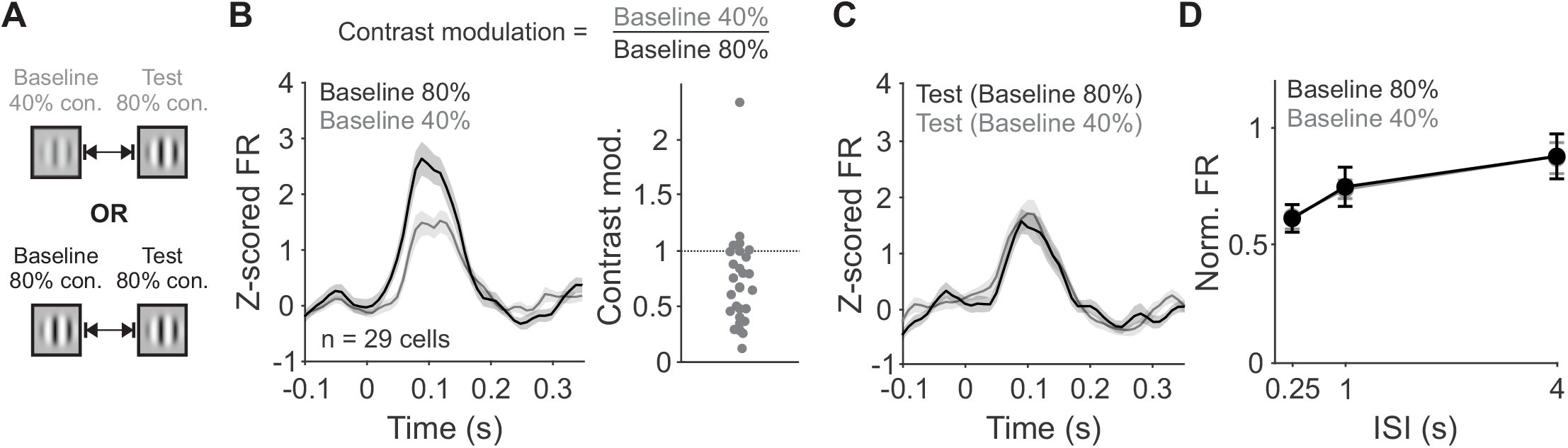
Effect of low contrast baseline stimulus on adaptation in L2/3, related to Figure 8. **A**. Schematic of visual stimulus. Baseline stimulus was either low (40%) or high (80%) contrast and test stimulus was always high contrast. **B**. Left: Z-scored PSTH of L2/3 units during high contrast (black) or low contrast (gray) baseline visual stimulus presentation. Right: Fractional change in peak firing rate during baseline stimulus for high versus low contrast. **C**. Z-scored PSTH during test visual stimulus presentation with baseline high (black) or low contrast (gray). **D**. Average normalized firing rate (test/baseline) with high or low contrast baseline stimulus. Test responses for both trial types was divided by the baseline response to high contrast (two-way ANOVA, effect of contrast, p = 0.45).

## References

1. Weber, A.I., Krishnamurthy, K., and Fairhall, A.L. (2019). Coding Principles in Adaptation. Annu. Rev. Vis. Sci. 5, 427–449. 10.1146/annurev-vision-091718-014818.

2. Barlow, H.B.H. (1961). Possible principles underlying the transformation of sensory messages. In Sensory Communication, pp. 217–234. 10.1080/15459620490885644.

3. Kohn, A. (2007). Visual adaptation: Physiology, mechanisms, and functional benefits. J. Neurophysiol. 97, 3155–3164. 10.1152/jn.00086.2007.

4. Vinje, W.E., and Gallant, J.L. (2000). Natural Vision Sparse Coding and Decorrelation in Primary Visual Cortex During Sparse Coding and Decorrelation in Primary Visual Cortex During Natural Vision. Science (80-.). 287, 1273–1276. 10.1126/science.287.5456.1273.

5. Simoncelli, E.P., and Olshausen, B.A. (2001). Natural Image Statistics and Neural Representation. Annu. Rev. Neurosci. 24, 1193–1216. 10.1146/annurev.neuro.24.1.1193.

6. Schwartz, O., Hsu, A., and Dayan, P. (2007). Space and time in visual context. Nat. Rev. Neurosci. 8, 522–535. 10.1038/nrn2155.

7. Parker, P.R.L., Abe, E.T.T., Leonard, E.S.P., Martins, D.M., and Niell, C.M. (2022). Joint coding of visual input and eye/head position in V1 of freely moving mice. Neuron, 1–10. 10.1016/j.neuron.2022.08.029.

8. Nigam, S., Milton, R., Pojoga, S., and Dragoi, V. (2023). Adaptive coding across visual features during free-viewing and fixation conditions. Nat. Commun. 14, 87. 10.1038/s41467-022-35656-w.

9. Latimer, K.W., Barbera, D., Sokoletsky, M., Awwad, B., Katz, Y., Nelken, I., Lampl, I., Fairhall, A.L., and Priebe, N.J. (2019). Multiple timescales account for adaptive responses across sensory cortices. J. Neurosci. 39, 10019–10033. 10.1523/JNEUROSCI.1642-19.2019.

10. Ulanovsky, N., Las, L., Farkas, D., and Nelken, I. (2004). Multiple time scales of adaptation in auditory cortex neurons. J. Neurosci. 24, 10440–10453. 10.1523/JNEUROSCI.1905-04.2004.

11. Zucker, R.S., and Regehr, W.G. (2002). Short-term synaptic plasticity. Annu. Rev. Physiol. 64, 355–405. 10.1146/annurev.physiol.64.092501.114547.

12. Varela, J.A., Sen, K., Gibson, J., Fost, J., Abbott, L.F., and Nelson, S.B. (1997). A quantitative description of short-term plasticity at excitatory synapses in layer 2/3 of rat primary visual cortex. J. Neurosci. 17, 7926–7940.

13. Abolafia, J.M., Vergara, R., Arnold, M.M., Reig, R., and Sanchez-Vives, M. V. (2011). Cortical auditory adaptation in the awake rat and the role of potassium currents. Cereb. Cortex 21, 977–990. 10.1093/cercor/bhq163.

14. Beck, O., Chistiakova, M., Obermayer, K., and Volgushev, M. (2005). Adaptation at synaptic connections to layer 2/3 pyramidal cells in rat visual cortex. J. Neurophysiol. 94, 363–376. 10.1152/jn.01287.2004.

15. Baccus, S.A., and Meister, M. (2002). Fast and slow contrast adaptation in retinal circuitry. Neuron 36, 909–919. 10.1016/S0896-6273(02)01050-4.

16. Patterson, C.A., Wissig, S.C., and Kohn, A. (2013). Distinct Effects of Brief and Prolonged Adaptation on Orientation Tuning in Primary Visual Cortex. J. Neurosci. 33, 532–543. 10.1523/JNEUROSCI.3345-12.2013.

17. Wolfe, J.M. (1984). Short test flashes produce large tilt aftereffects. Vision Res. 24, 1959–1964. 10.1016/0042-6989(84)90030-0.

18. Harris, J.P., and Calvert, J.E. (1989). Contrast, spatial frequency and test duration effects on the tilt aftereffect: Implications for underlying mechanisms. Vision Res. 29, 129–135. 10.1016/0042-6989(89)90179-X.

19. Jin, M., Beck, J.M., and Glickfeld, L.L. (2019). Neuronal adaptation reveals a suboptimal decoding of orientation tuned populations in the mouse visual cortex. J. Neurosci., 3172–18. 10.1523/JNEUROSCI.3172-18.2019.

20. Jin, M., and Glickfeld, L.L. (2020). Magnitude, time course, and specificity of rapid adaptation across mouse visual areas. J. Neurophysiol. 124, 245–258. 10.1152/jn.00758.2019.

21. Fritsche, M., Solomon, S.G., and de Lange, F.P. (2022). Brief Stimuli Cast a Persistent Long-Term Trace in Visual Cortex. J. Neurosci. 42, 1999–2010. 10.1523/jneurosci.1350-21.2021.

22. Sanchez-Vives, M. V, Nowak, L.G., and McCormick, D.A. (2000). Membrane mechanisms underlying contrast adaptation in cat area 17 in vivo. J Neurosci 20, 4267–4285. 20/11/4267 [pii].

23. Carandini, M., and Ferster, D. (1997). A Tonic Hyperpolarization Underlying Contrast A Tonic Hyperpolarization Underlying Contrast Adaptation in Cat Visual Cortex. 276, 949–952. 10.1126/science.276.5314.949.

24. Wehr, M., and Zador, A.M. (2005). Synaptic mechanisms of forward suppression in rat auditory cortex. Neuron 47, 437–445. 10.1016/j.neuron.2005.06.009.

25. Chung, S., Li, X., and Nelson, S.B. (2002). Short-term depression at thalamocortical synapses contributes to rapid adaptation of cortical sensory responses in vivo. Neuron 34, 437–446. 10.1016/S0896-6273(02)00659-1.

26. Natan, R.G., Briguglio, J.J., Mwilambwe-Tshilobo, L., Jones, S.I., Aizenberg, M., Goldberg, E.M., and Geffen, M.N. (2015). Complementary control of sensory adaptation by two types of cortical interneurons. Elife 4, 1–27. 10.7554/elife.09868.

27. Boudreau, C.E., and Ferster, D. (2005). Short-term depression in thalamocortical synapses of cat primary visual cortex. J. Neurosci. 25, 7179–7190. 10.1523/JNEUROSCI.1445-05.2005.

28. Hu, B., Garrett, M.E., Groblewski, P.A., Ollerenshaw, D.R., Shang, J., Roll, K., Manavi, S., Koch, C., Olsen, S.R., and Mihalas, S. (2021). Adaptation supports short-term memory in a visual change detection task. PLoS Comput. Biol. 17, 1–22. 10.1371/journal.pcbi.1009246.

29. Sohya, K., Kameyama, K., Yanagawa, Y., Obata, K., and Tsumoto, T. (2007). GABAergic neurons are less selective to stimulus orientation than excitatory neurons in layer II/III of visual cortex, as revealed by in vivo functional Ca2+ imaging in transgenic mice. J. Neurosci. 27, 2145–2149. 10.1523/JNEUROSCI.4641-06.2007.

30. Niell, C., and Stryker, M. (2008). Highly Selective Receptive Fields in Mouse Visual Cortex. J. Neurosci. 28, 7520–7536. 10.1523/JNEUROSCI.0623-08.2008.

31. Kerlin, A.M., Andermann, M.L., Berezovskii, V.K., and Reid, R.C. (2010). Broadly Tuned Response Properties of Diverse Inhibitory Neuron Subtypes in Mouse Visual Cortex. Neuron 67, 858–871. 10.1016/j.neuron.2010.08.002.

32. Hage, T.A., Bosma-Moody, A., Baker, C.A., Kratz, M.B., Campagnola, L., Jarsky, T., Zeng, H., and Murphy, G.J. (2022). Synaptic connectivity to L2/3 of primary visual cortex measured by two-photon optogenetic stimulation. Elife 11, 1–46. 10.7554/elife.71103.

33. Lefort, S., and Petersen, C.C.H. (2017). Layer-Dependent Short-Term Synaptic Plasticity between Excitatory Neurons in the C2 Barrel Column of Mouse Primary Somatosensory Cortex. Cereb. Cortex 27, 3869–3878. 10.1093/cercor/bhx094.

34. Mahn, M., Saraf-Sinik, I., Patil, P., Pulin, M., Bitton, E., Karalis, N., Bruentgens, F., Palgi, S., Gat, A., Dine, J., et al. (2021). Efficient optogenetic silencing of neurotransmitter release with a mosquito rhodopsin. Neuron 109, 1621–1635.e8. 10.1016/j.neuron.2021.03.013.

35. Chance, F.S., Nelson, S.B., and Abbott, L.F. (1998). Synaptic depression and the temporal response characteristics of V1 cells. J. Neurosci. 18, 4785–4799.

36. Abbott, L.F., Varela, J.A., Sen, K., and Nelson, S.B. (1997). Synaptic depression and cortical gain control. Science (80-.). 275, 220–224. 10.1126/science.275.5297.221.

37. Feldmeyer, D., Lübke, J., Silver, R.A., and Sakmann, B. (2002). Synaptic connections between layer 4 spiny neurone-layer 2/3 pyramidal cell pairs in juvenile rat barrel cortex: Physiology and anatomy of interlaminar signalling within a cortical column. J. Physiol. 538, 803–822. 10.1113/jphysiol.2001.012959.

38. Higley, M.J. (2006). Balanced Excitation and Inhibition Determine Spike Timing during Frequency Adaptation. J. Neurosci. 26, 448–457. 10.1523/JNEUROSCI.3506-05.2006.

39. Whitmire, C.J., and Stanley, G.B. (2016). Rapid Sensory Adaptation Redux: A Circuit Perspective. Neuron 92, 298–315. 10.1016/j.neuron.2016.09.046.

40. Heiss, J.E., Katz, Y., Ganmor, E., and Lampl, I. (2008). Shift in the balance between excitation and inhibition during sensory adaptation of S1 neurons. J. Neurosci. 28, 13320–13330. 10.1523/JNEUROSCI.2646-08.2008.

41. Carandini, M., Heeger, D.J., and Senn, W. (2002). A synaptic explanation of suppression in visual cortex. J. Neurosci. 22, 10053–10065. 10.1523/jneurosci.22-22-10053.2002.

42. Gabernet, L., Jadhav, S.P., Feldman, D.E., Carandini, M., and Scanziani, M. (2005). Somatosensory integration controlled by dynamic thalamocortical feed-forward inhibition. Neuron 48, 315–327. 10.1016/j.neuron.2005.09.022.

43. Crowder, N.A., Price, N.S.C., Hietanen, M.A., Dreher, B., Clifford, C.W.G., and Ibbotson, M.R. (2006). Relationship between contrast adaptation and orientation tuning in V1 and V2 of cat visual cortex. J. Neurophysiol. 95, 271–283. 10.1152/jn.00871.2005.

44. Borst, J.G.G. (2010). The low synaptic release probability in vivo. Trends Neurosci. 33, 259–266. 10.1016/j.tins.2010.03.003.

45. Chen, C., and Regehr, W.G. (2003). Presynaptic modulation of the retinogeniculate synapse. J. Neurosci. 23, 3130–3135. 10.1523/jneurosci.23-08-03130.2003.

46. Stratford, K.J., Tarczy-Hornoch, K., Martin, K.A.C., Bannister, N.J., and Jack, J.J.B. (1996). Excitatory synaptic inputs to spiny stellate cells in cat visual cortex. Nature 382, 258–261. 10.1038/382258a0.

47. Gil, Z., Connors, B.W., and Amitai, Y. (1997). Differential regulation of neocortical synapses by neuromodulators and activity. Neuron 19, 679–686. 10.1016/S0896-6273(00)80380-3.

48. Litvina, E.Y., and Chen, C. (2017). An evolving view of retinogeniculate transmission. Vis. Neurosci. 34. 10.1017/S0952523817000104.

49. Hirsch, J.A., Martinez, L.M., Alonso, J.M., Desai, K., Pillai, C., and Pierre, C. (2002). Synaptic physiology of the flow of information in the cat’s visual cortex in vivo. J. Physiol. 540, 335–350. 10.1113/jphysiol.2001.012777.

50. Nelson, S.B., and Turrigiano, G.G. (1998). Synaptic depression: A key player in the cortical balancing act. Nat. Neurosci. 1, 539–541. 10.1038/2775.

51. Seay, M.J., Natan, R.G., Geffen, M.N., and Buonomano, D. V. (2020). Differential short-term plasticity of PV and SST neurons accounts for adaptation and facilitation of cortical neurons to auditory tones. J. Neurosci. 40, 9224–9235. 10.1523/JNEUROSCI.0686-20.2020.

52. Hu, H., and Agmon, A. (2016). Differential excitation of distally versus proximally targeting cortical interneurons by unitary thalamocortical bursts. J. Neurosci. 36, 6906–6916. 10.1523/JNEUROSCI.0739-16.2016.

53. Wright, N.C., Borden, P.Y., Liew, Y.J., Bolus, M.F., Stoy, W.M., Forest, C.R., and Stanley, G.B. (2021). Rapid Cortical Adaptation and the Role of Thalamic Synchrony during Wakefulness. J. Neurosci. 41, 5421–5439. 10.1523/JNEUROSCI.3018-20.2021.

54. Yarden, T.S., Mizrahi, A., and Nelken, I. (2022). Context-Dependent Inhibitory Control of Stimulus-Specific Adaptation Context-Dependent Inhibitory Control of Stimulus-Specific Adaptation Abbreviated Title: 4 Inhibitory Control of Stimulus-Specific Adaptation. 42, 4629–4651.

55. Yavorska, I., and Wehr, M. (2016). Somatostatin-expressing inhibitory interneurons in cortical circuits. Front. Neural Circuits 10, 1–18. 10.3389/fncir.2016.00076.

56. Beierlein, M., Gibson, J.R., and Connors, B.W. (2003). Two Dynamically Distinct Inhibitory Networks in Layer 4 of the Neocortex. J. Neurophysiol. 90, 2987–3000. 10.1152/jn.00283.2003.

57. Heintz, T.G., Hinojosa, A.J., Dominiak, S.E., and Lagnado, L. (2022). Opposite forms of adaptation in mouse visual cortex are controlled by distinct inhibitory microcircuits. Nat. Commun. 13, 1–14. 10.1038/s41467-022-28635-8.

58. Phillips, E.A.K., Schreiner, C.E., and Hasenstaub, A.R. (2017). Cortical Interneurons Differentially Regulate the Effects of Acoustic Context. Cell Rep. 20, 771–778. 10.1016/j.celrep.2017.07.001.

59. Hamm, J.P., and Yuste, R. (2016). Somatostatin Interneurons Control a Key Component of Mismatch Negativity in Mouse Visual Cortex. Cell Rep. 16, 597–604. 10.1016/j.celrep.2016.06.037.

60. Varela, J.A., Song, S., Turrigiano, G.G., and Nelson, S.B. (1999). Differential depression at excitatory and inhibitory synapses in visual cortex. J. Neurosci. 19, 4293–4304. 10.1523/jneurosci.19-11-04293.1999.

61. Kapfer, C., Glickfeld, L.L., Atallah, B. V, and Scanziani, M. (2007). Supralinear increase of recurrent inhibition during sparse activity in the somatosensory cortex. Nat. Neurosci. 10, 743–753. 10.1016/j.immuni.2010.12.017.Two-stage.

62. Reyes, A., Lujan, R., Rozov, A., Burnashev, N., Somogyi, P., and Sakmann, B. (1998). Target-cell-specific facilitation and depression in neocortical circuits. Nat. Neurosci. 1, 279–284. 10.1038/1092.

63. Pala, A., and Petersen, C.C.H. (2015). InVivo Measurement of Cell-Type-Specific Synaptic Connectivity and Synaptic Transmission in Layer 2/3 Mouse Barrel Cortex. Neuron 85, 68–75. 10.1016/j.neuron.2014.11.025.

64. Scholl, B., Thomas, C.I., Ryan, M.A., Kamasawa, N., and Fitzpatrick, D. (2020). Cortical response selectivity derives from strength in numbers of synapses. Nature 12. 10.1038/s41586-020-03044-3.

65. Petersen, C.C.H., and Crochet, S. (2013). Synaptic Computation and Sensory Processing in Neocortical Layer 2/3. Neuron 78, 28–48. 10.1016/j.neuron.2013.03.020.

66. Tsodyks, M. V., and Markram, H. (1997). The neural code between neocortical pyramidal neurons depends on neurotransmitter release probability. Proc. Natl. Acad. Sci. U. S. A. 94, 719–723. 10.1073/pnas.94.2.719.

67. Chance, F.S., and Abbott, L.F. (2001). Input-specific adaptation in complex cells through synaptic depression. Neurocomputing 38–40, 141–146. 10.1016/S0925-2312(01)00550-1.

68. Foley, J.M., and Boynton, G.M. (1993). Forward pattern masking and adaptation: Effects of duration, interstimulus interval, contrast, and spatial and temporal frequency. Vision Res. 33, 959–980. 10.1016/0042-6989(93)90079-C.

69. Nagel, K.I., Hong, E.J., and Wilson, R.I. (2015). Synaptic and circuit mechanisms promoting broadband transmission of olfactory stimulus dynamics. Nat. Neurosci. 18, 56–65. 10.1038/nn.3895.

70. Horwitz, G.D. (2020). Temporal information loss in the macaque early visual system 10.1371/journal.pbio.3000570.

71. Reinhold, K., Lien, A.D., and Scanziani, M. (2015). Distinct recurrent versus afferent dynamics in cortical visual processing. Nat. Neurosci. 18, 1789–1797. 10.1038/nn.4153.

72. Douglas, R.J., and Martin, K.A.C. (2004). Neuronal Circuits of the Neocortex. Annu. Rev. Neurosci. 27, 419–451. 10.1146/annurev.neuro.27.070203.144152.

73. Cheetham, C.E.J., and Fox, K. (2010). Presynaptic development at L4 to L2/3 excitatory synapses follows different time courses in visual and somatosensory cortex. J. Neurosci. 30, 12566–12571. 10.1523/JNEUROSCI.2544-10.2010.

74. Voelcker, B., Pancholi, R., and Peron, S. (2022). Transformation of primary sensory cortical representations from layer 4 to layer 2. Nat. Commun. 13, 5484. 10.1038/s41467-022-33249-1.

75. Castro-Alamancos, M.A. (2004). Absence of Rapid Sensory Adaptation in Neocortex during Information Processing States. Neuron 41, 455–464. 10.1016/S0896-6273(03)00853-5.

76. Edelstein, A.D., Tsuchida, M.A., Amodaj, N., Pinkard, H., Vale, R.D., and Stuurman, N. (2014). Advanced methods of microscope control using μManager software. J. Biol. Methods 1, e10. 10.14440/jbm.2014.36.

77. Sanzeni, A., Akitake, B., Goldbach, H.C., Leedy, C.E., Brunel, N., and Histed, M.H. (2020). Inhibition stabilization is a widespread property of cortical networks. Elife 9, 1–39. 10.7554/eLife.54875.

